# SARS-CoV-2 spike protein induces abnormal inflammatory blood clots neutralized by fibrin immunotherapy

**DOI:** 10.1101/2021.10.12.464152

**Authors:** Jae Kyu Ryu, Elif G. Sozmen, Karuna Dixit, Mauricio Montano, Yusuke Matsui, Yixin Liu, Ekram Helmy, Thomas J. Deerinck, Zhaoqi Yan, Renaud Schuck, Rosa Meza Acevedo, Collin M. Spencer, Reuben Thomas, Alexander R. Pico, Scott S. Zamvil, Kara L. Lynch, Mark H. Ellisman, Warner C. Greene, Katerina Akassoglou

## Abstract

Blood clots are a central feature of coronavirus disease-2019 (COVID-19) and can culminate in pulmonary embolism, stroke, and sudden death. However, it is not known how abnormal blood clots form in COVID-19 or why they occur even in asymptomatic and convalescent patients. Here we report that the Spike protein from severe acute respiratory syndrome coronavirus 2 (SARS-CoV-2) binds to the blood coagulation factor fibrinogen and induces structurally abnormal blood clots with heightened proinflammatory activity. SARS-CoV-2 Spike virions enhanced fibrin-mediated microglia activation and induced fibrinogen-dependent lung pathology. COVID-19 patients had fibrin autoantibodies that persisted long after acute infection. Monoclonal antibody 5B8, targeting the cryptic inflammatory fibrin epitope, inhibited thromboinflammation. Our results reveal a procoagulant role for the SARS-CoV-2 Spike and propose fibrin-targeting interventions as a treatment for thromboinflammation in COVID-19.

**One-Sentence Summary:** SARS-CoV-2 spike induces structurally abnormal blood clots and thromboinflammation neutralized by a fibrin-targeting antibody.

---

Persistent life-threatening thrombotic events are a hallmark of COVID-19. Aberrant clots form in multiple organs causing significant morbidity and mortality in COVID-19 patients (*1, 2*). The high incidence of clotting complications has been attributed to disease severity, inflammation and subsequent hypercoagulable state (*3*). However, the clinical picture is puzzling because of disproportionate rates of thrombotic events and abnormal clot properties not observed in other inflammatory conditions, such as severe sepsis or different viral respiratory illnesses (*4-7*). Intriguingly, abnormal clotting is not limited to acutely-ill COVID-19 patients. Pulmonary emboli, stroke and sudden death also occur in young COVID-19 patients with asymptomatic infections or mild respiratory symptoms (*8*). Persistent clotting pathology is prevalent in post-acute sequelae of SARS-CoV-2 infection (PASC, Long COVID) (*2, 7, 9*).

The central structural component of blood clots, and a key regulator of inflammation in disease, is insoluble fibrin, which is derived from the blood coagulation factor fibrinogen and is deposited in tissues at sites of vascular damage (*10, 11*). Hypercoagulability in COVID-19 is associated with inflammation and the formation of fibrin clots resistant to degradation despite adequate anticoagulation (*3-5*). Extensive fibrin deposits are detected locally in inflamed lung and brain tissues from COVID-19 patients, sometimes without evidence of direct viral infection at autopsy (*1, 8, 12-14*). The high prevalence of thrombotic events with these unique hypercoagulability features suggests an as yet unknown mechanism of abnormal blood clot formation in COVID-19. We set out to determine how blood clots form in COVID-19 and to identify therapies to combat the deleterious effects of abnormal coagulation occurring in acute and convalescent stages of disease.

Since hypercoagulability in COVID-19 patients has features distinct from those of other inflammatory diseases, we hypothesized that SARS-CoV-2 directly affects the structural and functional properties of blood clots. Incubation of SARS-CoV-2 recombinant trimeric spike protein (Spike) with healthy donor plasma increased fibrin polymerization (Fig. 1A). Spike strikingly altered the fibrin clot structure resulting in thinner fibers with a rough appearance and increased clot density as shown by scanning electron microscopy (SEM) (Fig. 1B, fig. S1), identifying direct effects of SARS-CoV-2 Spike on fibrin clot architecture. Consistent with these structural changes, a solid-phase binding assay revealed binding of Spike to both fibrinogen and fibrin (K_d_ 5.3 µM and 0.4 µM, respectively) (Fig. 1C). Fibrinogen immunoprecipitated with full-length recombinant trimeric Spike, and studies with deletion mutants identified an interaction with the S2 domain of Spike (Fig. 1D, fig. S2). Fibrinogen is a 340 kDa protein consisting of three pairs of polypeptide chains Aα, Bβ, and γ (*10*). To identify Spike binding regions in fibrinogen, we generated a custom fibrinogen peptide array consisting of 390 15-mer peptides overlapping by 11 amino acids and spanning the fibrinogen Aα, Bβ, and γ chains (Fig. 1E). Hybridization with His-tagged trimeric Spike identified three binding sites in the Bβ and γ fibrinogen chains, namely Bβ119-129, γ_163-181_ and γ_364-395_ fibrinogen peptides (Fig. 1E). The Bβ_119-129_ peptide contains cleavage sites for the fibrinolytic serine protease plasmin (*15*). Spike bound to the γ_364-395_ peptide, which encompasses the γ_377-395_ cryptic fibrinogen binding site to complement receptor 3 that activates innate immune responses (*11, 16*). Spike also bound to the γ_163-181_ peptide, whose function is unknown. Mapping of the Spike binding peptides onto the crystal structure of fibrinogen revealed proximity of the γ_163-181_ and γ_377-395_ peptides, suggesting that a 3D conformational epitope in the carboxy-terminal γ-chain of fibrinogen (γC domain) is involved in fibrinogen binding to Spike.

**Fig. 1.**
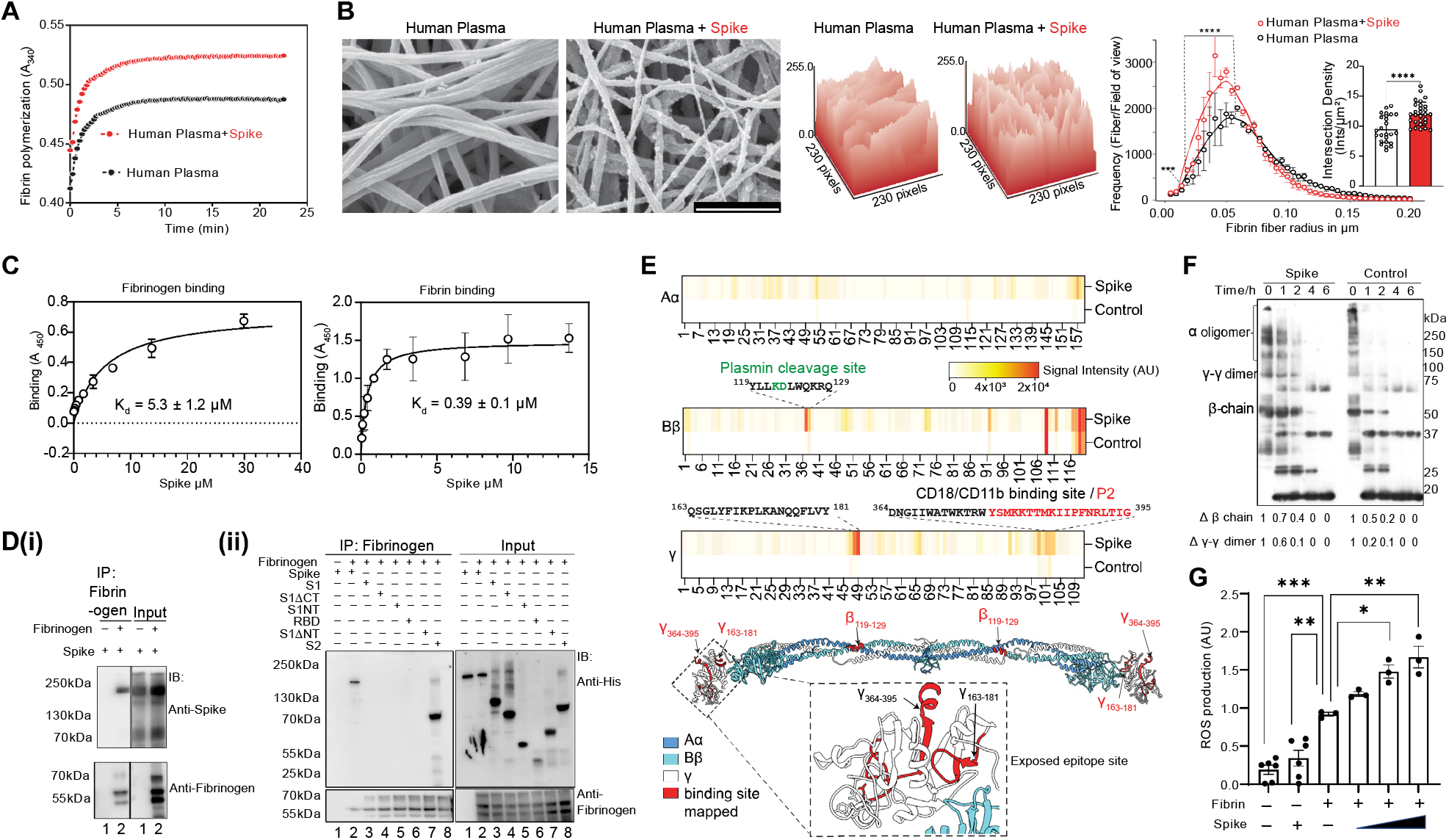
Characterization of the interaction between SARS-CoV-2 Spike and fibrinogen. (**A**) Turbidity assay of fibrin polymerization in healthy human donor plasma in the presence or absence of Spike protein. Data are representative of four independent experiments with similar results. (**B**) SEM images of fibrin clots in healthy human donor plasma or in the presence of Spike protein. Scale bar, 1 µm. Topographic visualization of fibrin fiber surface in SEM images. Quantification of fibrin fiber radius and intersection density. Data are from three independent experiments (mean ± s.e.m.). ^****^*P* < 0.0001, ^***^*P* < 0.001 (multiple testing Holm procedure and two-tailed Mann-Whitney test). (**C**) ELISA of the binding of recombinant SARS-CoV-2 Spike protein (Spike) to fibrinogen or fibrin presented as absorbance at 450 nm (A_450_), plus the dissociation constants (K_d_). Representative binding curvefits are shown from two independent experiments performed in duplicates (mean ± s.e.m.). (**D)**. Immunoprecipitation of fibrinogen with His-tagged recombinant trimeric SARS-CoV-2 Spike protein produced in CHO cells (i) or monomeric SARS-CoV-2 Spike produced in *E*.*coli* (ii) blotted with anti-spike, anti-His or anti-fibrinogen. Representative immunoblots from three independent experiments are shown. (**E**) Peptide array mapping with immobilized peptides of fibrinogen chains Aα, Bβ, and γ blotted with Spike protein. Heatmap of signal intensity showing binding sites (red-orange) indicated by their aa sequences on chains Bβ (β_119-129_) and γ (γ_163-181_ and γ_364-395_). Color key indicates fluorescence intensities signal values from low (white) to high (red). Crystal structure of fibrinogen (PDB:3GHG) showing the three mapped peptides β_119-129_, γ_163-181_ and γ_364-395_ (red). Structural proximity of the γ_163-181_ and γ_364-395_ peptides indicating a 3D conformational epitope (inset). (**F**) Immunoblot of fibrin degradation after 0, 1, 2, 4 and 6 h of plasmin digestion. Data are from five (timepoints 0, 2 and 4 h) or three (timepoints 1 and 6 h) independent experiments (mean ± s.e.m.). Representative immunoblot is shown. (**G**) Quantification of ROS production detected with dihydroethidum in unstimulated BMDMs or stimulated for 24 h with fibrin in the presence of Spike protein. Data are from three independent experiments (mean ± s.e.m.). ^*^*P* < 0.05; ^**^*P* < 0.01; ^***^*P* < 0.001 (one-way analysis of variance (ANOVA) with Tukey’s multiple comparisons test).

Since Spike binds to fibrinogen sites that regulate plasmin cleavage and binding to complement receptor 3, we tested whether the binding interferes with the degradation and inflammatory properties of fibrin. Incubation of Spike with fibrin delayed plasmin degradation of both the β-chain and the γ-γ dimer (Fig. 1F), suggesting that Spike delays fibrinolysis. This finding is consistent with dense fibrin clots composed of thin fibers we identified and the presence of fibrinolysis-resistant blood clots in COVID-19 patients (*5*). Dense fibrin clots with thin fibers resistant to lysis are also observed in thromboembolic diseases (*17*).

Fibrin is deposited locally at sites of vascular damage and is a potent proinflammatory activator and a key inducer of oxidative stress (*11, 18*). Strikingly, Spike increased fibrin-induced release of reactive oxygen species (ROS) in a concentration-dependent manner in bone marrow-derived macrophages (BMDMs), while Spike alone did not have an effect (Fig. 1G). These results suggest a role for Spike as an enhancer of fibrin-induced inflammation at sites of vascular damage. Overall, these results reveal an unanticipated role for SARS-CoV-2 Spike as a fibrinogen binding protein that alone accelerates the formation of abnormal clots with altered structure and increased inflammatory activity.

In COVID-19 patients, fibrin is deposited in the air spaces and lung parenchyma and is associated with inflammation (*8*). We developed an experimental platform to study the interplay between fibrin and SARS-CoV-2 Spike *in vivo* by injecting mice with HIV virions pseudotyped with SARS-CoV-2 trimeric Spike glycoprotein (Spike PVs) (fig. S3), enabling the study of the *in vivo* effects of Spike independent of active viral replication. Intravenous administration of Spike PVs in wild-type (WT) mice induced extensive fibrin deposition in the lung (Fig. 2A). Double immunofluorescence staining for fibrin and Spike PVs revealed strong overlap of Spike and fibrin deposits (Fig. 2B, movie S1, S2). Fibrin deposition was associated with activated endothelium in the lung, and gene expression analysis revealed increased expression of endothelial and inflammatory markers in Spike PV-injected mice compared to mice injected with control BALD PVs (Fig. 2C, fig. S4, tables S1, S2), consistent with findings of SARS-CoV-2 toxicity to endothelial cells (*19*). Fibrin activates macrophages and induces oxidative stress through nicotinamide adenine dinucleotide phosphate (NADPH) oxidase (*11, 18*), which is linked to severe disease and thrombotic events in COVID-19 patients (*20*). In WT mice, Spike-PVs activated macrophages and increased expression of the gp-91-phox subunit of NADPH oxidase in the lung of WT mice indicating the generation of an oxidative stress response (Fig. 2D). In contrast, control BALD PVs or PVs expressing the Env protein from the HIV-1(HIV-1 PVs) did not induce these effects (Fig. 2D), suggesting that lung pathology was specific for SARS-CoV-2 Spike. Mice genetically-deficient in fibrinogen (*Fgα*^*–/–*^ mice), which express all other blood proteins except fibrinogen and are protected from autoimmune and inflammatory conditions (*11*), did not exhibit lung pathology following Spike PV challenge (Fig. 2E, fig. S5). These results reveal a Spike– fibrinogen-dependent mechanism of clot formation that generates strong inflammatory and oxidative stress responses.

**Fig. 2.**
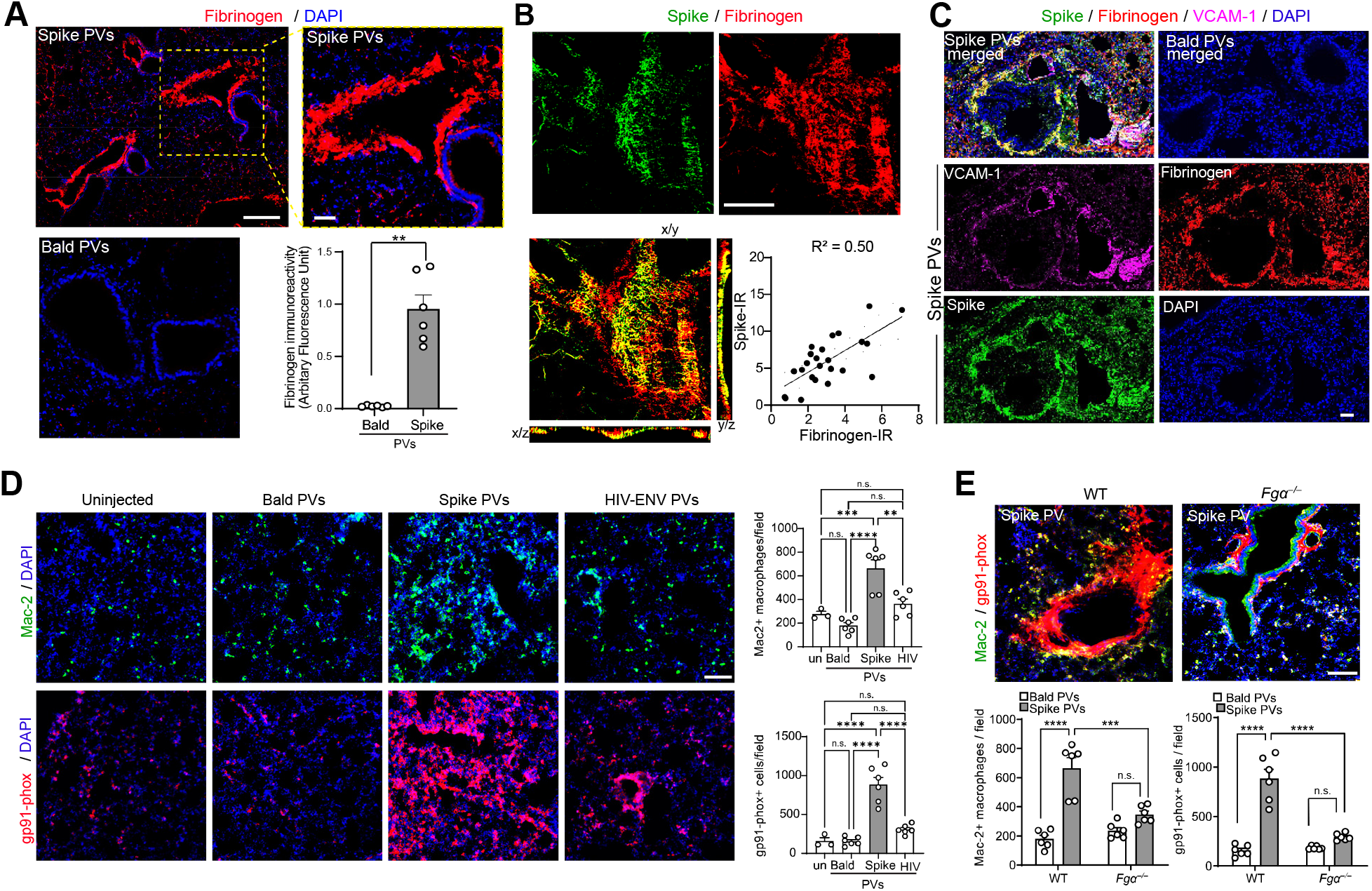
Fibrinogen-dependent SARS CoV-2 Spike lung patholog. (**A**) Microscopy of mouse lung sections obtained 24 h after injection of BALD or Spike PVs and stained with anti-fibrin(ogen) antibody. Nuclei are stained with 4′,6-diamidino-2-phenylindole (DAPI; blue). Scale bars, 200 µm (top left panel) and 50 µm (top right panel). Data are from *n* = 6 mice per group (mean ± s.e.m.). ^**^*P* < 0.01 (two-tailed Mann-Whitney test). (**B**) Confocal microscopy of immunofluorescence double-stained mouse lung sections obtained 24 h after injection of Spike PVs, showing immunoreactivity of Spike (green) and fibrinogen (red); orthogonal views of the y/z and x/z planes show the localization of fibrinogen and Spike. Scale bar, 50 μm. Representative images from three mice are shown. Scatter plot shows a positive correlation of fibrinogen and Spike immunoreactivity in 24 images from *n* = 3 mice, R^2^ = 0.496, *P* = 0.0001, Pearson correlation. (**C**) Confocal microscopy of mouse lung sections 24 h after Spike PV injection showing VCAM-1, fibrinogen, and Spike immunoreactivity. Representative images from three mice are shown. (**D**) Microscopy of lung sections from mice 24 h after injection with BALD, Spike, or HIV-ENV PVs 24 h and from uninjected healthy controls (un), showing immunoreactivity of Mac-2 (green) and gp91-phox (red). Scale bars, 100 μm. Data are from *n* = 6 mice (BALD, SARS-CoV-2 Spike, and HIV-ENV PVs) and *n* = 3 mice (uninjected controls) (mean ± s.e.m.). ^**^*P* < 0.01, ^***^*P* < 0.001, ^****^*P* < 0.0001 (one-way ANOVA with Tukey’s multiple comparisons test). (**E**) Microscopy of the lung of control and *Fgα*^*–/–*^ mice showing Mac- 2 and gp91-phox immunoreactivity after Spike PV injection. Scale bar, 100 µm. Data are from *n* = 6 mice per group (mean ± s.e.m.). ^***^*P* < 0.001, ^****^*P* < 0.0001 (two-way ANOVA with Tukey’s multiple comparisons test).

Fibrinogen is causally linked to the activation of macrophages and microglia in autoimmune and inflammatory diseases in the brain and periphery (*11, 21*). Fibrin is a driver of microglia-induced cognitive dysfunction (*22*) and is associated with perivascular-activated microglia and macrophages in brains of COVID-19 patients even without signs of infection (*12*). Stereotactic co-injection of Spike PVs and fibrinogen into the brains of WT mice, a model of fibrinogen-induced encephalomyelitis (*21*), significantly increased fibrin-induced microglia activation (Fig. 3A), suggesting that Spike enhances the inflammatory function of fibrin *in vivo*. Furthermore, like recombinant Spike (Figs. 1D, G), Spike PVs co-immunoprecipitated with fibrinogen and increased fibrin-induced oxidative stress in BMDMs (figs. S6, S7). Conversion of fibrinogen to fibrin exposes the cryptic inflammatory γ_377–395_ epitope in the fibrinogen γ-chain. Genetic or pharmacologic targeting of this epitope has potent therapeutic effects in autoimmune and inflammatory diseases (*11, 18, 23, 24*). Using alanine scanning mutagenesis, we found that Spike interacts with aa 386-394 in the C-terminus of the γ_377–395_ epitope (Fig. 3B, fig. S8). Genetic targeting of the fibrin γ_377–395_ epitope in *Fgg*^*γ390-396A*^ mice, in which fibrinogen retains normal clotting function but lacks the γ_390-396_-binding motif, rescued from macrophage activation and oxidative stress in the lung after Spike PV administration (Fig. 3C). Since γ_377–395_ is a binding site for both Spike (this study) and complement receptor (*16, 18, 23, 24*), inhibition of this epitope may reduce their interactions with fibrin. Future studies will be required to characterize the biophysical properties of fibrin binding to Spike and to complement receptor 3 and their relative contributions to thromboinflammation. These findings reveal a previously unknown interaction between SARS-CoV-2 Spike protein and fibrin γ_377–395_ epitope that promotes innate immune activation.

**Fig. 3.**
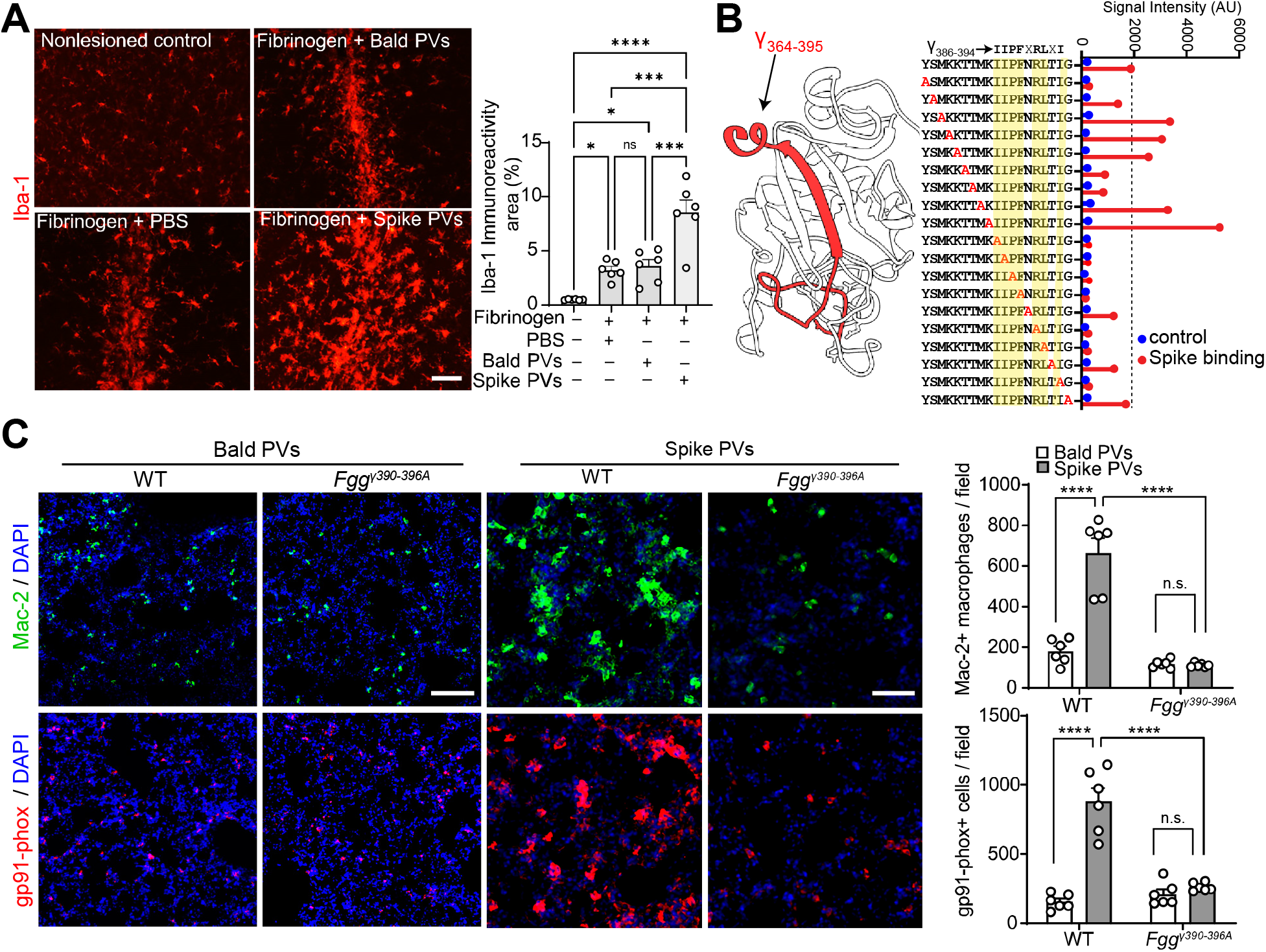
The fibrin γ_377–395_ cryptic epitope is required for innate immune activation by SARS CoV-2 Spike. (**A**) Microscopy of brain sections from control or stereotaxic co-injection of fibrinogen with PBS, BALD, or Spike PVs, showing Iba-1 immunoreactivity. Scale bar, 50 µm. Data are from *n* = 6 mice per group (mean ± s.e.m.). ^*^*P* < 0.05, ^***^*P* < 0.001, ^****^*P* < 0.0001 (one-way ANOVA with Tukey’s multiple comparisons test). n.s., not significant. (**B**) Structural map of the carboxyl-terminal γ-chain (white) showing the mapped Spike-binding epitope γ_364-395_ (red). Alanine scanning mutagenesis of peptide γ_377-395_ blotted with His-tagged Spike protein. Signal intensity bar graph of the binding of Spike to sequential Ala substituted peptides (red). Residues with low signal intensity upon Ala substitution required for binding are highlighted yellow. (**C**) Microscopy of lung sections from WT and *Fgg*^*γ390-396A*^ mice 24 h after injection of BALD or Spike PVs showing immunoreactivity for Mac-2 and gp91-phox. Scale bars, 50 μm. Data are from *n* = 6 mice per group (mean ± s.e.m.). ^***^*P* < 0.001, ^****^*P* < 0.0001 (two-way ANOVA with Tukey’s multiple comparisons test).

A surge of autoantibody production against diverse immune targets have been detected in COVID-19 patients (*25*). To determine whether COVID-19 patients develop autoantibodies against abnormal blood clots, we tested autoantibody responses to fibrin. Autoantibodies against fibrin epitopes would be potentially missed by the inherent limitations of phage and yeast library screens to produce post-translationally modified insoluble fibrin polymer. To overcome this challenge, we developed a fibrin autoantibody discovery platform optimized for screening patient samples. We tested longitudinally collected serum samples ranging from acute to convalescent disease stages from 54 COVID-19 asymptomatic, mild, and severe disease patients requiring admission to the intensive care units (table S3). Fibrin autoantibodies were abundant in all three groups of COVID-19 patients and persisted during the convalescent stage, but were scarce in healthy donor controls or in subjects with non-COVID respiratory illnesses (Fig. 4A, B).

**Fig. 4.**
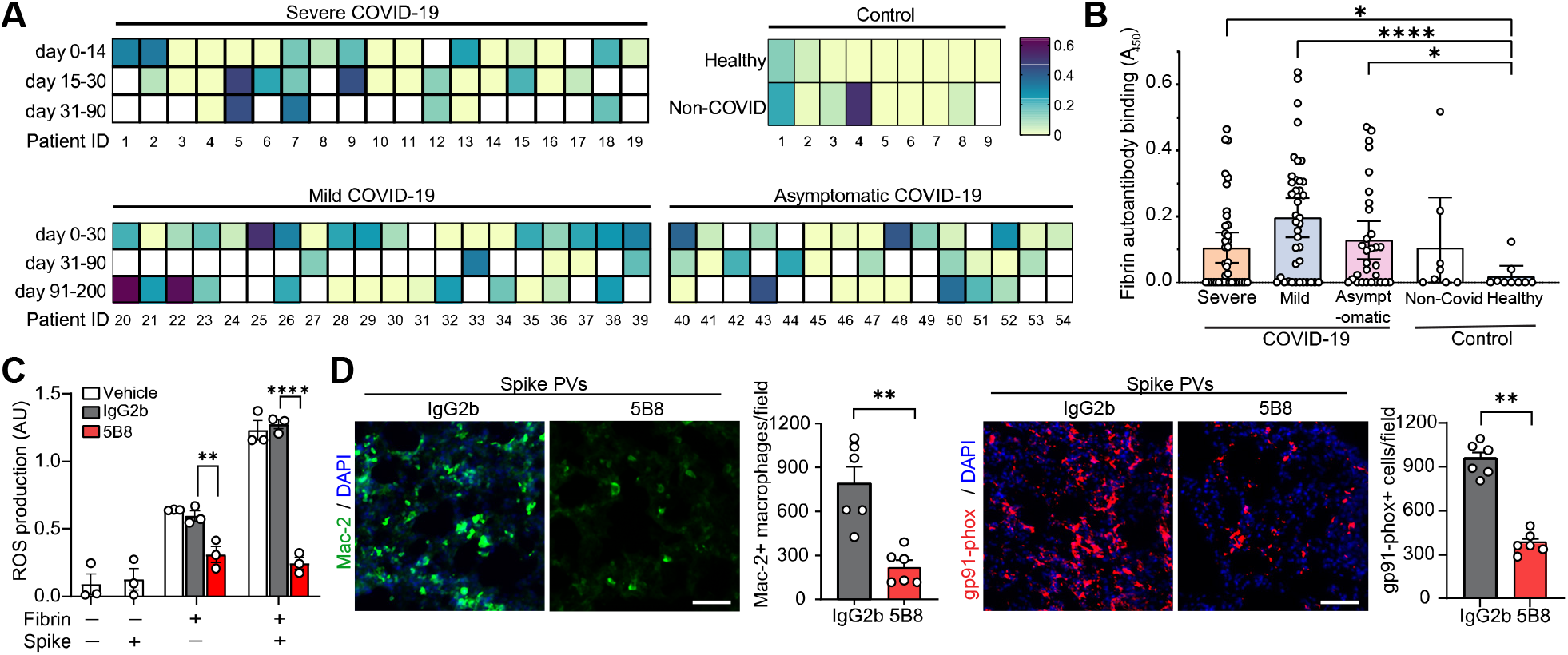
Fibrin-targeting immunotherapy protects from SARS-CoV-2 Spike thromboinflammation. (**A**) Heatmap of fibrin autoantibody ELISA from longitudinal serum collections across at least one time point from patients with severe (*n* = 19), mild (*n* = 20) and asymptomatic (*n* = 15) COVID-19 patients vs. 9 healthy controls and 8 controls with non-COVID respiratory illnesses. White boxes indicate unavailable samples for specific time points. (**B**) Whole group sample comparison of fibrin autoantibody levels in samples from patients with severe (*n* = 38), mild (*n* = 39), or asymptomatic (*n* = 31) COVID-19, vs. 9 healthy controls and 8 controls with non-COVID-19 respiratory illness. Values are mean ± s.e.m. ^*^*P =* 0.0207, ^****^*P* < 0.0001, ^*^*P* = 0.0121, n.s. = nonsignificant (Welch’s ANOVA with Dunnett’s multiple comparisons test). (**C**) ROS production in unstimulated BMDM or stimulated for 24 h with Spike or/and fibrin after 5B8 or IgG2b treatment. Data are from three independent experiments (mean ± s.e.m.). (**D**) Microscopy of lung sections from WT mice injected with Spike PVs, and either 5B8 (30 mg/kg) or IgG2b (30 mg/kg), showing Mac-2 and gp91-phox immunoreactivity. Scale bar, 50 μm. Data are from *n* = 6 mice per group (mean ± s.e.m.). ^**^*P* < 0.01 (two-tailed Mann-Whitney test).

Blockade of the thromboinflammation cascade following Spike and fibrinogen/fibrin interaction is an attractive therapeutic target. Based on our genetic rescue results implicating a causal role for the γ_377–395_ epitope, we tested the effects of 5B8, a monoclonal antibody generated against the fibrin γ_377–395_ epitope (*18*). This selective antibody-based approach suppresses fibrin-induced inflammation without altering normal hemostasis (*18*). 5B8 rescued the enhanced inflammatory effects induced by Spike in fibrin-treated BMDMs (Fig. 4C), suggesting that pharmacologic blockade of the fibrin γ_377–395_ epitope inhibits the deleterious effects of SARS-CoV-2 Spike as an enhancer of thromboinflammation. Strikingly, the 5B8 antibody reduced macrophage activation and oxidative stress in the lungs of Spike PV-treated WT mice compared to isotype IgG2b-treated controls (Fig. 4D). Collectively, these results identify anti-fibrin autoimmune responses in COVID-19 patients and demonstrate a potent protective effect of fibrin-targeting immunotherapy against thromboinflammation.

In summary, we find that SARS-CoV-2 Spike protein enhances the formation of highly inflammatory clots that are neutralized by a fibrin-targeting monoclonal antibody. Our data shed new light on the enigmatic coagulopathy found in COVID-19 revealing a causal role for fibrinogen in thromboinflammation – even independent of active viral replication. The high incidence of clotting complications in COVID-19 has been attributed to systemic inflammation (*3*), vascular damage including abnormal levels of circulating coagulation proteins (*1, 26*), genetic susceptibility to tissue factor and complement genes (*27*), and prothrombotic autoantibodies (*28*). Our findings now show that coagulopathy is not merely a consequence of inflammation. Rather, the interaction of SARS-CoV-2 Spike with fibrinogen and fibrin results in abnormal blood clot formation that in turn drives inflammation. Identification of SARS-CoV-2 Spike protein as a fibrinogen binding partner provides a mechanistic basis for the formation of abnormal clots with enhanced inflammatory properties. This mechanism might be in play at sites of local fibrin deposition and microvascular injury perpetuating a hypercoagulable and inflammatory state as reported in COVID-19 patients (*14*) that could be critical during acute infection, as well as in PASC (*2, 5, 7*). Fibrin is locally deposited in brain and other organs of COVID-19 patients. Thus, fibrin immunotherapy may represent a novel strategy for reducing thromboinflammation in systemic and neurologic manifestations of COVID-19. Since anti-fibrin antibody 5B8 has protective effects (this study) and protective autoantibodies have been reported in COVID-19 patients (*29*), some fibrin autoantibodies may be protective against thromboinflammation. Whether the human fibrin autoantibodies in COVID-19 reported here have potential divergent functions, or associate with clinical characteristics, or have biomarker value, warrants further investigation. Furthermore, the fibrin-Spike coagulation and inflammation assays we describe can serve as a discovery platform for identifying new therapeutics for COVID-19. Targeting fibrin may be an efficacious therapy to suppress thromboinflammation in both acute COVID-19 and PASC patients.

## Acknowledgments

We thank Deepak Srivastava and Melanie Ott for critical reading of the manuscript, Florian Krammer for sharing the plasmid for mammalian expression of recombinant Spike, Andrew S. Mendiola, Min-Gyoung Shin, Eunbi L. Ryu and the Gladstone Flow Cytometry Core for technical assistance, and Stephen Ordway for editorial assistance.

## Funding

The Roddenberry Foundation (WCG and KA)

National Institutes of Health grants NS120055, R24GM137200 (MHE)

National Institutes of Health grant R35 NS097976 (KA)

National Institutes of Health T32 AI007334 (EGS)

National Science Foundation NSF2014862-UTA20-000890 (MHE)

James B. Pendleton Charitable Trust (WCG)

Edward and Pearl Fein (KA)

Simon Family Trust (KA)

## Author contributions

Conceptualization: KA, JKR, WCG

Methodology: JKR, EGS, MM, MHE, TJD, WCG, KA

Software: RS

Validation: EGS, MM, KD, YM, YL, EH, MHE, WCG, KA

Formal analysis: RT, ARP, JKR, EGS, KD, YL, ZY, RS

Investigation: JKR, EGS, KD, MM, YM, YL, EH, TJD, ZY, KLY, RMA, MHE

Resources: CMS, KLY, SSZ, MHE, WCG, KA

Data curation: JKR, EGS, KD, MHE, RS, RT, ARP, WCG, KA

Writing-original draft: KA, EGS, JKR, KD

Writing-Review & Editing: all authors, Supervision: KA, WCG, MHE

Project Administration: KA, WCG

Funding acquisition: KA, WCG, MHE, EGS

## Competing interests

KA is a co-founder, scientific advisor, and board director of Therini Bio. KA is the inventor of the 5B8 patent. KA, JKR, MM, and WCG. are inventors on a patent application for 5B8 use in COVID-19. KA, JKR, MM, EGS and WCG are inventors on a patent application related to Spike-induced thromboinflammation model. KA and JKR are inventors on fibrin assay patents. Their interests are managed in accordance with their respective institutions’ conflict of interest policy.

## Data and materials availability

Expression plasmid pCAGGS seq nCoV19 of soluble trimeric spike deposited by Florian Krammer is freely available from BEI Resources. Distribution of Spike PVs, mice, and antibodies to non-profit investigators will be under a Material Transfer Agreement (MTA). All data are available in the manuscript or the supplementary materials.

