## Supplementary Materials for "SARS-CoV-2 spike protein induces abnormal inflammatory blood clots neutralized by fibrin immunotherapy"

**This PDF file includes:**

Materials and Methods

Figs. S1 to S8

Tables S1 to S3

Captions for Movies S1, S2

**Other Supplementary Material for this manuscript includes the following:**

Movies S1, S2

### Materials and Methods

#### Subjects and specimens

The COVID-19 infection cohort utilized remnant serum samples from routine clinical laboratory testing at Zuckerberg San Francisco General Hospital (ZSFG). All patients had positive results by SARS-CoV-2 real-time polymerase chain reaction (RT-PCR) in nasopharyngeal swabs between March and July 2020. Clinical data were extracted from electronic health records and included demographic information, major co-morbidities, patient-reported symptom onset date, symptoms, and indicators of disease severity. COVID-19 was classified as severe in 19 patients (admitted to intensive care unit), mild in 20 patients (admitted to hospital or managed as outpatients), and as asymptomatic in 15 patients (not hospitalized, no symptoms). The criteria for ICU admission at the hospital remained the same throughout the course of the study. Non-COVID respiratory illness control (n=8) remnant samples were from febrile patients with upper respiratory symptoms who were SARS-CoV-2 RT-PCR negative and positive for another respiratory virus (BioFire® FilmArray® Respiratory 2 Panel, Salt Lake City, UT). The protocol for ZSFG remnant specimen collection from patients with suspected COVID-19 infection (IRB #20-30387) was approved by the UCSF Institutional Review Board. The committee judged that written consent was not required for use of remnant specimens. Whole blood from healthy control were obtained from subjects enrolled at the University of California, San Francisco (UCSF) for a study that predates the COVID-19 outbreak. Human subjects research was approved by the UCSF Institutional Review Board (IRB# 10-00650). All subjects provided written informed consent before participation in this study. 10 ml whole blood was collected from each healthy control subject into Vacutainer tubes without anticoagulant, and blood was allowed to coagulate for 30 minutes. Tubes were centrifuged at 1300 *x g* for 30 min. Serum layer was harvested, aliquoted, and preserved at -80°C.

#### Animals

C57BL/6 mice were purchased from the Jackson Laboratory. $Fg\alpha^{-/-}$ (*1*) and $Fgg^{\gamma390-396A}$ mice (*2*) were obtained from Dr. Jay Degen (University of Cincinnati, OH, USA). Mice were housed under a 12:12 light/dark cycle, 55% ± 5% relative humidity, and a temperature of 20 ± 2 °C with access to standard laboratory chow and water *ad libitum*. They were housed in social groups of a maximum of 5 mice in standard mouse housing cages and bedding. All single -housed mice were provided with cage enrichment (a cardboard or hard-plastic house-like hiding place and tissue paper). For husbandry, one male and one female were housed together with a maximum of one litter was permitted. Mice were weaned at postnatal day 21. Male mice were used for all experiments. All animal procedures were performed under the guidelines set by the Institutional Animal Care and Use Committee at the University of California, San Francisco.

**SARS-CoV-2 recombinant trimeric spike protein production in a mammalian expression system**

The plasmid vector pCAGGS containing the SARS Coronavirus 2,Wuhan-Hu-1 ectodomain Spike glycoprotein gene with a deletion of the polybasic cleavage site (RRAR to A), two stabilizing mutations (K986P and V987P), a C-terminal thrombin cleavage site, T4 foldon trimerization domain, and a hexahistidine tag (6xHis) was obtained from BEI Resources (deposited by the laboratory of Dr. Florian Krammer) *(3)*. Recombinant Spike protein was produced by Celltheon (Union City, CA). Briefly, CHO cells were transiently transfected with the plasmid and harvested at >70% viability. Spike protein was obtained by centrifugation and sterile filtration, purified by $Ni^{2+}$-NTA affinity chromatography, and eluted in phosphate-buffered saline (PBS) containing imidazole. Fractions containing eluted recombinant spike protein were then buffer exchanged into 1x PBS and was further purified by size-exclusion chromatography using a Superdex 200 column.

**Fibrin polymerization assay**

Fibrin polymerization was measured by turbidity analysis as described (*4*). In brief, pooled healthy donor citrated human plasma (Innovative Research) was diluted to 1:3 in 20 mM HEPES and 137 mM NaCl (pH 7.4). Recombinant trimeric spike protein was buffer exchanged into 20 mM HEPES, pH 7.4, 137 mM NaCl using Amicon concentrators (100 kDa cut-off) prior to plasma incubation. 50 µl of plasma dilution was incubated with 50 µl recombinant trimeric spike protein at 25 °C for 15 min. Recombinant trimeric spike protein was freshly thawed without freezing and thaw cycles. Clotting was initiated by 0.25 U/ml thrombin (Sigma-Aldrich) and 20 mM $CaCl_2$. Final concentrations were 1:12 plasma, 0.75 µM Spike, 0.25 U/ml thrombin, 20 mM $CaCl_2$. Turbidity was measured at 340 nm every 15 sec for 30 min in a SpectraMax M5 microplate reader (Molecular Devices) with SoftMax Pro 5.2 software (Phoenix Technologies).

**Scanning electron microscopy (SEM) of fibrin clots**

Healthy donor citrated human plasma was diluted 1:3 in 20 mM HEPES buffer, pH 7.4; 15 µl of the diluted plasma was mixed with 15 µl of recombinant trimeric spike protein that was buffer exchanged into 20 mM HEPES and 137 mM NaCl using Amicon concentrators (100 kDA cut-off) prior to addition to plasma. Low concentration of NaCl was used to maintain spike solubility and stability. Then, 25 µl of this mixture was pipetted onto 5 mm x 5 mm silicon wafers (Ted Pella) and incubated for 15 min at 37 °C in a humidified tissue culture incubator. Next, 25 µl of a solution of $CaCl_2$ and thrombin in 20 mM HEPES was added in the center of the wafer and allowed to polymerize at 25 °C for 2 h. Final concentrations were plasma 1:12, 0.9 µM Spike, 0.25 U/ml thrombin, 20 mM $CaCl_2$. Buffer was used instead of spike for control condition. Clots on wafers were placed on ice, washed twice for 10 min each with ice-cold electron microscopy grade 0.1 M cacodylate buffer, pH 7.4, and fixed in cold electron microscopy grade 2% glutaraldehyde (Electron Microscopy Sciences). Samples were rinsed three times for 5 min each in Millipore-filtered, double-distilled water; dehydrated in an ethanol series (20%, 50%, 70%, 90%, 100%, 100% for 2 min each); and critical point dried with $CO_2$. Samples were sputter coated with a thin layer of gold-palladium and imaged with a Zeiss Merlin field-emission scanning electron microscope at 3.0 keV and a secondary electron detector.

SEM imaging and image acquisition were carried out blinded to test conditions. 4000X images were captured across the sample, then were converted to 8-bit with NIH ImageJ (v. 1.50). After pixel to micron scaling, each image was cropped into two to three fields of view (FOV) (8×8 µm) with NIH DiameterJ as described (*5*). Briefly, the best segmentation algorithm was pre-selected based on side by side comparison of images before quantification. Surface plot plug-in ImageJ was used to generate topographical maps of SEM images. The Mixed Segmentation (M1-through M3 options) built in DiameterJ Segment provided the most accurate representation of the fibers to be quantified. The same segmentation method and variant was used across all test conditions and images. Each segmented image was manually edited with ImageJ to ensure complete representation of segmented fibers. Edited images were batch processed with DiameterJ 1-108 (orientation analysis not selected). Fiber radius and intersection densities were collated from each batch. Data from 8–10 FOVs per sample was generated for group analysis.

**Fibrinogen and fibrin coated ELISA Plates**

Fibrinogen and fibrin coated plates were prepared as described (*4*). Briefly, human plasminogen-free fibrinogen (EMD Millipore) was used after IgG depletion using a Pierce albumin/IgG removal kit (Thermo Fisher Scientific). IgG-depleted human plasminogen-free fibrinogen was further diluted to 25 μg/ml by adding 20 mM HEPES buffer, pH 7.4 for coating fibrinogen plates or 20 mM HEPES buffer pH 7.4 with 1U/ml thrombin (Sigma-Aldrich) and 7 mM $CaCl_2$ for fibrin coated plates. Coating was performed for 1.5 h at 37 °C using 96-well MaxiSorp plates (Thermo Fisher Scientific) and fibrin-coated plates were dried at 37 °C overnight as described (*4*).

**Recombinant SARS-CoV-2 Spike protein binding on fibrin or fibrinogen**

Fibrin- or fibrinogen-coated 96-well plates were washed with wash buffer (1×PBS + 0.05% Tween-20), and incubated with blocking buffer consisting of wash buffer with 5% bovine serum albumin (BSA) (Omnipure, Fisher) for 1 h at 25 °C. Serial dilutions of recombinant spike were made in binding buffer (wash buffer containing 0.5% BSA). Recombinant trimeric spike protein was added to the wells and incubated for 2 h at 37 °C. After washing five times with binding buffer, rabbit polyclonal anti-6x His tag antibody (1:1000, abcam, ab137839) was added to the plates and incubated for 1 h at 25 °C. Following washing, goat anti-rabbit IgG H&L (conjugated with horse radish peroxidase, HRP) (1:1000, abcam, ab205718) in wash buffer was added for 1 h at 25 °C. After the final wash, the HRP substrate 3,3′,5,5′-tetramethybenzidine (TMB; Sigma-Aldrich) was added into the wells. The reaction was quenched by adding 1N hydrochloric acid, and absorption was measured at 450 nm. Non-linear regression curves were analyzed using Prism 9 software to calculate $K_d$ values using one site binding model.

**Fibrinogen peptide array and spike binding epitope mapping**

A custom PepStar$^{TM}$ Multiwell fibrinogen Peptide array that comprises a purified synthetic peptide library containing 390 15-mer peptides representing overlapping peptide scans (15/11) of the α, β, and γ fibrinogen chains (UniProt IDs: FIBA P02671, FIBB P02675, FIBG P02679) was generated by JPT Peptide Technologies (Berlin, Germany). The arrays were hybridized with Recombinant-His tagged trimeric Spike protein (1 μg/ml in blocking buffer) for 1 h at 30 °C. The His-tag peptide (AGHHHHHH) was co-immobilized on the peptide microarray slides as an assay control. Microarrays were incubated for 1 h at 30 °C with fluorescently labeled anti-6xHis monoclonal antibody (Alexa 647, Invitrogen, MA1-135-A647) diluted to 1 μg/ml in blocking buffer and dried. Before each step, microarrays were washed with washing buffer, 50 mM TBS-buffer including 0.1% Tween20, pH 7.2. The assay buffer was LowCross buffer (Candor Bioscience). The slides were washed, dried, and scanned with a high-resolution laser scanner at 635 nm to obtain fluorescence intensity profiles. The images were quantified to yield a mean pixel value for each peptide. All incubations were done in 1 day. To assess nonspecific binding to the peptides and assay performance, a control incubation with secondary antibody only (no sample present) was done in parallel on each slide. The resulting images were analyzed and quantified with spot-recognition software (GenePix, Molecular Devices). For each spot, the mean signal intensity was extracted (between 0 and 65535 arbitrary units). To visualize the results, heatmap diagrams representing all peptides immobilized on the microarray and containing all signal values were computed; fluorescence intensities were color-coded from white (no binding), to yellow (medium binding), to red (strong binding). Binding peptides were further mapped onto the 3D crystal structure of fibrinogen (PDB ID: 3GHG) with UCSF Chimera (*6*).

**Peptide alanine scanning**

Alanine scanning experiments were performed with PepStar™ Multiwell microarrays containing 60 peptides representing Alanine substitutions of each residue on peptide YSMKKTTMKIIPFNRLTIG ($\gamma_{377\text{-}395}$) by JPT Peptide Technologies. Human full-length IgG and His-tagged peptides were co-immobilized on the peptide microarray slides as assay controls. His-tagged spike protein was applied at five concentrations (from 10 μg/ml to 0.001 μg/ml) and incubated for 1 h at 30 °C. Two fluorescently labeled secondary antibodies specific to the His tag were added separately and left to react for 1 h. After washing and drying, the slides were scanned with a high-resolution laser scanner at 635 nm to obtain the fluorescence intensity profiles, and the images were quantified to yield a mean pixel value for each peptide. Control incubations with each secondary antibody only (no sample present) were performed in parallel on the slide to assess nonspecific binding to the peptides and assay performance. The data was analyzed with respect to the original peptide. A higher signal after alanine substitution indicated that a residue was not involved in binding to spike protein, if; a lower signal indicated that a residue was important for binding to spike protein.

**Plasmin digestion of fibrin clots**

Before clotting, 3 μM fibrinogen was incubated with 9 μM recombinant trimeric spike protein at 37 °C for 1 h in 20 mM HEPES, pH 7.4, 137 mM NaCl, 5 mM $CaCl_2$. Thrombin was added to the mixture at a final concentration of 1.5 U/ml. Fibrin clots were allowed to form in Eppendorf tubes over a 2-h incubation at 37 °C. Then, 5 µl of 100 µg/ml plasmin (Millipore) was added to each tube on top of the clot. All samples were incubated at 37 °C for 0, 1, 2, 4, and 6 h; digestion was quenched by adding sodium dodecyl sulfate–polyacrylamide gel electrophoresis (SDS-PAGE loading buffer with reducing regent. Samples were heated at 85 °C for 20 min, and aliquots (equivalent to 100 ng fibrinogen) were separated by SDS-PAGE on 4–12% Bis-Tris gels, transferred to PVDF membranes, and analyzed for anti-human fibrinogen by western blot. Band intensities of each protein species (i.e., γ-γ dimer, β-chain) were analyzed with Image J and normalized to corresponding bands at the 0 h time point.

**ROS detection**

BMDM cell culture and ROS detection using 5 μM DHE (Invitrogen) were performed as described (*4, 7*). Briefly, cells were plated on 96-well black μ-clear-bottomed microtiter plates (Greiner Bio-One) pre-coated with 12.5 μg/ml fibrin with or without recombinant trimeric spike protein (0.168, 1.68, and 3.36 μM). For fibrin inhibition, 5B8 or IgG2b (each 20 μg/ml) (MPC-11, BioXCell) was added in fibrin with or without 3.36 μM recombinant trimeric spike protein-coated wells 2 h before plating of cells. Cells were incubated on fibrin for 24 h and DHE fluorescence was detected at 518 nm/605 nm with a SpectraMax M5 microplate reader. Since macrophage activation can be influenced by cell culture conditions, heat-inactivated fetal bovine serum and macrophage colony-stimulating factor were batch tested as described (*7*).

**Production of recombinant monomeric Spike and monomeric Spike deletion mutants in *E. coli***

A plasmid expressing full-length Spike (amino acids (aa) 1–1273) of SARS-CoV-2, Wuhan-Hu-1 (GenBank: MN908947) with a C-terminal 6xHis was generated by amplifying the Spike coding sequence and inserting it into pET-21a(+) (Novagen) at *BamH*I/*Xho*I sites. Plasmids expressing six Spike mutants—S1 (aa 1–685), S1ΔCT (aa 1–541), S1NT (aa 1–318), receptor binding domain (aa 319–541), ST1ΔNT (aa 319–685), and S2 (aa 686–1273)—were generated with a PCR-based method and properly mutated primers. The expression plasmids were transformed into *E. coli* Rosetta 2(DE3) pLysS competent cells (Novagen) and cultured in 500-ml flasks containing 100 ml of LB + ampicillin and chloramphenicol on a shaking incubator (250 rpm, 37 °C) to optical density of 0.4 at 600 ng. The cultures were induced with 1 mM IPTG (Thermo Fisher Scientific), incubated for 4 h, and centrifuged (6000 *g*, 4 °C) for 15 min. The supernatant was removed, and the pellets were frozen at –80 °C overnight. Spike produced in *E.coli* was only used for immunoprecipitation assays.

**Immunoprecipitation assay**

For co-immunoprecipitation to test interaction of fibrinogen with Spike protein (His-tagged recombinant trimeric spike protein or monomeric Spike protein produced in *E. coli*), the Pierce co-immunoprecipitation kit (Thermo Fisher Scientific) protocol was used with an original immunoprecipitation/lysis buffer and modifications. For lysis, the frozen cell pellets were solubilized in 800 µl of immunoprecipitation/lysis buffer (50 mM Tris, pH 8.0, 5% glycerol, 1% NP-40, 100 mM NaCl) supplemented with 100 µg/ml lysozyme (Sigma-Aldrich), 100× EDTA-free Halt protease inhibitor (Thermo Fisher Scientific), and 250 U/µl benzonase nuclease (Sigma-Aldrich). *E. coli* cells were lysed by two rounds of sonication at 30 Hz for 30 sec each until the sample was no longer viscous. After further mixing for 20 min in a rotator, the lysate was cleared by centrifugation at 10,000 *g* for 10 min, warmed to 37 °C, mixed with 25 µg of fibrinogen, incubated for 1 h, applied to beads incubated with 10 µg of anti-fibrinogen sheep antibody (SAFG-AP, Enzyme Research Laboratories), and incubated for 1 h at 37 °C. All washes were done essentially as described in the kit. The bound proteins were eluted in 60 µl of EB solution provided in the kit and neutralized with 1/10 volume of 1 M Tris, pH 9.0. Washing buffer and EB solution were warmed to 37 °C in advance.

The eluted proteins were separated by SDS-PAGE on 4–12% gels, transferred to PVDF membranes (Invitrogen), and incubated with rabbit anti-His antibody (1:1000, Cell Signaling, 2365S) and sheep anti-fibrinogen antibody (1:1000, Enzyme Research Laboratories, SAFG-AP) and then with HRP-conjugated anti-rabbit (111-035-144, Jackson ImmunoResearch; 1:10,000) and anti-sheep (HAF016, R&D Systems; 1:5000) secondary antibodies. Protein bands were detected with Immobilon Forte Western HRP substrate (Sigma-Aldrich) and the ChemiDoc imaging system (Bio-Rad).

**Production of Spike pseudotyped virions (PVs)**

For production of HIV virions pseudotyped with SARS-CoV-2 trimeric Spike glycoprotein (Spike PVs), 293T cells ($3.75\times10^6$) were plated in a T175 flask and transfected 24 h later with 90 µg of polyethyleneimine (PEI), 30 µg of HIV-1 NL-4-3 Δ Env eGFP (NIH AIDS Reagent Program), or 3.5 µg of pCAGGS SARS-CoV-2 trimeric Spike glycoprotein (NR52310, BEI Resources) in a total of 10 ml of Opti-MEM medium (Invitrogen), using PEI transfection reagent (Sigma). These pseudotyped virions do not carry the genetic material of the SARS-CoV-2 virus other than Spike, which does not bind efficiently to the murine ACE2 receptor, enabling the study of the in vivo effects of Spike to be studied in the absence of viral infection. The next day, the medium was replaced with DMEM10 complete medium, and the cells were incubated at 37 ºC in 5% $CO_2$ for 48 h. The supernatant was then harvested, filtered with 0.22-µm Steriflip filters (EMD, Millipore), and ultracentrifuged at 25,000 rpm for 1.5 h at 4 ºC. The concentrated supernatant was removed, the pellets (viral particles) were resuspended in cold 1× PBS containing 1% fetal bovine serum, and aliquots were stored at –80 ºC in a biosecurity level 3 laboratory. For production of control viral particles not expressing the Spike glycoprotein (BALD), the same procedure was used but with the omission of the pCAGGS SARS-CoV-2 Spike vector transfection. HIV ENV pseudotyped viral particles were also produced with the same procedure, using an HIV89.6 ENV dual tropic (X4 and R5) expression vector (NIH AIDS Reagent Program) instead of the Spike expression vector.

**In vivo administration of SARS-CoV-2 Spike PVs**

Mice were anaesthetized with isoflurane and placed on electric heating pad. Spike pseudotyped or BALD PVs (control) (100 µl) were slowly injected into the retro-orbital plexus with a BD 0.3-ml insulin syringe attached to a 29-gauge needle. After 3 min, the needle was slowly withdrawn, and the mice were allowed to recover. Since the activity of PVs can be influenced by freeze/thaw cycles, all experiments were done with virions that had been freshly thawed and kept at 37 °C. Refrozen virion samples were not used. SARS-CoV-2 Spike PVs were administered in 12- to 15-week-old male C57BL/6, $Fg\alpha^{-/-}$, and $Fgg^{\gamma390\text{–}396A}$ mice. Experiments in $Fg\alpha^{-/-}$ mice were done blinded to the mouse genotype. Experiments in $Fgg^{\gamma390\text{–}396A}$ mice were done blinded to the mouse genotype and the type of virions. Mice were randomly assigned to treatment groups in a blinded manner. Genotype and treatment assignment were revealed after image quantification.

**5B8 treatment**

For pharmacological treatment after SARS-CoV-2 Spike PV administration, anti-fibrin antibody 5B8 (*4*) or an isotype-matched IgG2b (MPC-11, BioXCell) control were administered by retro-orbital injection at 30 mg/kg 15 min before injection of Spike PVs to WT mice as described above. Mice were sacrificed at 24 h for histological analysis. Experimenters were blinded to treatment. Treatment assignments were revealed after histologic analysis and image quantification.

**Immunohistochemistry**

Lung tissues were cut with a cryostat into 30-μm-thick frozen sections for free-floating immunostaining. The following antibodies were used: mouse anti-SARS-CoV-2 (COVID-19) Spike antibody (1A9, GeneTex; 1:100), rat anti-mouse CD106 (VCAM-1, catalog no. 553849, BD Pharmingen; 1:100), and rabbit polyclonal anti-fibrinogen (gift from Dr. Jay Degen; 1:500). The tissue sections were washed in PBS and incubated in a blocking and permeabilization buffer consisting of PBS supplemented with 0.2% Triton X-100 and 5% BSA for 1 h at 25 °C. For mouse primary antibodies, sections were incubated in M.O.M. mouse IgG blocking reagent diluted in PBS containing 0.2% Triton X-100, and 5% BSA and then with M.O.M. diluent for 5 min at room temperature (M.O.M. (Mouse on Mouse) Immunodetection Kits, Vector Laboratories). Sections were rinsed twice with PBS containing 0.1% Triton X-100 and incubated overnight with primary antibodies at 4 °C. All tissue sections were washed with PBS containing 0.1% Triton X-100 and incubated with the following secondary antibodies: goat anti-rabbit Alexa Fluor 488 (1:1000, Thermo Fisher Scientific, A-11008), goat anti-mouse Alexa Fluor 568 (A-110041, Thermo Fisher Scientific; 1:1000), or goat anti-rat Alexa Fluor 647 (A-21247, Thermo Fisher Scientific; 1:1000), and stained with DAPI. Sections were mounted on frosted microscopic slides (Thermo Fisher Scientific), covered with glass coverslips, sealed with ProLong Diamond Antifade Mounting reagent (Thermo Fisher Scientific), and kept at 4 °C until imaging. To assess nonspecific antibody binding, negative control sections were incubated with Isotype-matched, nonspecific mouse, rat, and rabbit antibodies (catalog nos. 08-6599, 31933, and 08-6199, respectively, Thermo Fisher Scientific.

**Confocal microscopy**

Tissue sections were imaged with a laser-scanning confocal microscope (FLUOVIEW FV3000RS "Snow Leopard"), a 60× oil-immersion UPLSAPO objective (NA = 1.35), and FV31S-SW software, v 2.3.2.169 (Olympus). Individual channels were captured sequentially with a 405-nm laser and a 430/70 spectral detector for DAPI, a 488-nm laser and a 500/40 spectral detector for Alexa Fluor 488, a 561-nm laser and a 570/620 high-sensitivity detector for Alexa Fluor 568, and a 650-nm laser and a 650/750 high-sensitivity detector (Olympus TruSpectral detector technology) for Alexa Fluor 647. Captured images were processed with Fiji 2.1.0/Image J 1.53c.

**Image analysis**

Immunostained cells were counted with Jupyter Notebook in Python 3. Briefly, an arbitrary threshold was manually set and used for all images in the dataset. The total number of cells per image was estimated with the function "peak_local_max" from the open source "skimage" Python image processing library, which returns the coordinates and number of local peaks in an image (https://scikit-image.org/docs/dev/api/skimage.feature.html#skimage.feature.peak_local_max). Fibrinogen immunoreactivity was quantified with Fiji (ImageJ) as described (*8*). Python image processing was used to colocalize fibrinogen and Spike protein in lung tissues. Briefly, a Jupyter Notebook was written to estimate the amount of fluorescent signal overlap between Spike and fibrinogen in lung tissues. The "Ostu" filter from the "skimage" Python image processing library was used to threshold each image labeled with Spike and fibrinogen (https://scikit-image.org/docs/0.13.x/api/skimage.filters.html#skimage.filters.threshold_otsu). After thresholding, each set of images was compared, and pixels were compartmentalized in 4 categories: Spike and fibrinogen overlap, Spike signal only, fibrinogen signal only, and no signal. In each image, the total number of pixels in an image and the number of pixels with signal for Spike only, fibrinogen only, or both were computed for 24 images.

**Gene expression analysis**

SARS-CoV-2 Spike PVs were administered in male C57BL6/J mice as described above. 24 h after injection, mice were perfused with PBS following isolation of the lungs. A small piece of tissue from each lobe of the lung was dissected, combined, and immediately homogenized in a 1.5-ml Eppendorf tube with buffer RLT (Qiagen) and a pestle (catalog no. 749521-1500, Kimble Chase) on ice. The homogenate was further processed with QIAshredder (Qiagen), and RNA samples were extracted with the RNeasy Mini Kit (Qiagen). RNA concentration was measured with a Nanodrop spectrophotometer (catalog no. 840-274200, Thermo Scientific) and the integrity was determined with an Agilent Bioanalyzer. RNA samples were sent to the Core Center for Musculoskeletal Biology and Medicine at UCSF, and the gene expression of Mouse Immunology Panel (Codeset: NS_Immunology_Mm_C2269) was determined with a NanoString nCounter machine.

**Network analysis**

Network analysis was performed as described (*7*). The STRING database (*9, 10*) was queried for interactions among the 51 significantly upregulated genes (*P* value <0.05) after injection of Spike PVs relative to expression after injection of BALD PVs. A subset of 43 genes was connected by high-confidence interactions (score >0.8) and visualized with Cytoscape (*11*). Degree was calculated with the built-in Analyzer tool and mapped to node size, while the log2FoldChange was mapped to node fill color with a gradient over the full range of values (0.074–0.723). The network will be made available at NDEx under doi:10.18119/N9WK60.

**Stereotactic injection of fibrinogen and Spike**

Fibrinogen was stereotactically injected into the brain as described (*12*). Mice were anesthetized with isoflurane and placed in a stereotaxic apparatus (Kopf Instruments). Alexa Fluor 488 human fibrinogen (Thermo Fisher Scientific) was dissolved in 0.1 M sodium bicarbonate (pH 8.3) at 25 °C to a concentration of 1.5 mg/ml as described (*13*), mixed with Spike PVs, BALD PVs, or PBS control (1:1 ratio), and incubated at 37 ºC for 15 min; 1.5 µl of the mixture was stereotactically injected at 0.3 µl /min with a 10-µl Hamilton syringe and a 33-gauge needle into the corpus callosum of C57Bl/6 mice (*12*). The mice anesthetized with avertin and transcardially perfused with 4% paraformaldehyde in PBS. The brains were removed, postfixed overnight at 4 ºC in 4% paraformaldehyde in PBS, processed with 30% sucrose in PBS, cut into coronal sections 30 µm thick, washed in PBS, and incubated for 10 min with DAPI (Thermo Fisher Scientific; 1:1000) in PBS, and processed for immunohistochemical staining with rabbit anti-Iba-1 (catalog no. 019-19741, Wako; 1:1,000) as described (*7*). Images were acquired with an Axioplan II epifluorescence microscope (Zeiss) and Plan-Neofluar objectives (10 × 0.3 NA). Images of similar anatomical locations were quantified with NIH ImageJ (v. 1.50) by an observer blinded to experimental conditions. Images were acquired and quantified in a blinded manner. Treatment assignment were revealed after image quantification.

**Fibrin autoantibody detection in COVID-19 patient sera**

This experiment was carried out in the BSL-3 facility. Fibrin-coated plates were washed with 1× PBS and blocked with 4 mg/ml mouse serum solution (Molecular Biosciences, dissolved in 5% BSA in PBS) for 1 h at 25 °C. Wells were washed three times for 5 min each with wash buffer (0.05% Tween in 1× PBS). Frozen human serum samples were stored at –80 °C with minimal freeze and thaw except for aliquoting purposes until the screen. Once thawed on ice, each sample was gently mixed by pipetting, diluted 1:2 in sample dilution buffer (0.4 mg/ml mouse serum, 0.5% BSA, 0.05% Tween-20 in 1× PBS). Samples were plated in duplicate, incubated for 2 h at 37 °C on fibrin-coated plates. Wells were washed in wash buffer five times for 5 min each. Fibrin bound human IgG was detected by incubation with mouse anti-human IgG-HRP (1:5000, Invitrogen) for 1 h at 25 °C in the dark. After thorough washes in wash buffer, HRP substrate (TMB, Millipore) was added, and after adequate colorimetric development wells were neutralized with 1N hydrochloric acid. All samples shown were screened at the same time. Plates were read immediately at 450 nm. Individual well reads were corrected by subtracting secondary background signal.

**Statistical analysis**

All values are reported as mean ± s.e.m. Unless stated otherwise, $P$ values were calculated with one- or two-way analysis of variance followed Tukey's post hoc test for multiple comparisons or two-tailed Mann–Whitney test for non-normally distributed pairs in GraphPad Prism software 6.0. Sample sizes were determined by prior studies rather than statistical approaches. All mice survived until the end of the study, and all of the data was analyzed. For in vivo studies with $Fg\alpha^{-/-}$ mice, only mice, not virions, were randomized and coded for group assignment and data collection. For $Fgg^{\gamma390-396A}$ mice, both mice and virions were blindly assigned to experimental groups. For the antibody treatment, 5B8 and IgG2b were blindly assigned to experiment groups. For the NanoString experiment, virions were blindly assigned to experimental groups. All histological analysis and quantification were done in blinded fashion. No data were excluded. Biochemical studies of the binding of fibrinogen to Spike or Spike PVs were performed in the Akassoglou lab and independently validated in the Greene lab.

For quantification of the fibrin clots by scanning electron microscopy data, at each radius, we estimated the difference in log odds of detecting fibers (among all the views in a given image) with the chosen radius under Spike versus control conditions across all images (log odds ratio). The log odds ratio at each radius was estimated with generalized linear mixed effects models, with the family argument set to binomial and implemented in glmer function in the lme4 package in R (*14*), in which the image source for the observations is modeled as a random effect. *P* values were corrected for multiple testing with the Holm procedure (*15*). For co-localization of fibrin and Spike, the ratio of the odds that a pixel with signal for Spike would also have signal for fibrinogen to the odds that a pixel with Spike would not have signal for fibrinogen was estimated for each image with Fisher's exact test. For gene expression analysis from NanoString data, the normalized gene expression data from 4 replicates under each of Spike or BALD PV conditions were log 2 transformed. Up- or down-regulation of expression for each gene was estimated with a one-sided two-sample Welch *t* test. The resulting *P* values were corrected for multiple testing with the Benjamini-Hochberg procedure (*16*). All analyses were done in a given number of sections, or cells per lung tissue imaged *in vivo*, per mouse, and the mean ± s.e.m. was calculated for the reported number (n) of mice per group.

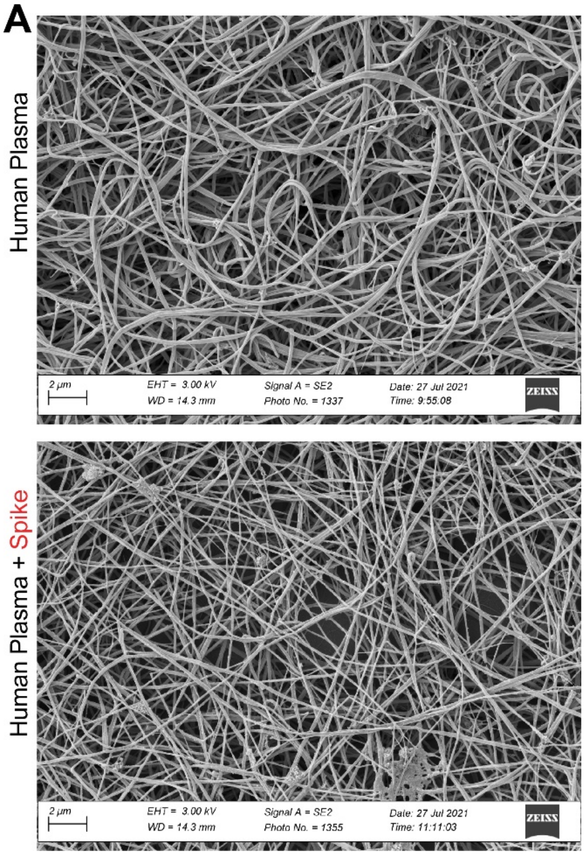


B

| radius (micron) | Log odds ratio | std.error | z.value | p value | p adjust |
|---|---|---|---|---|---|
| 0.0045 | -0.1662 | 0.1380 | -1.2042 | 2.2852E-01 | 4.5703E-01 |
| 0.0090 | 0.1813 | 0.0709 | 2.5582 | 1.0521E-02 | 3.6304E-02 |
| 0.0135 | 0.2579 | 0.0696 | 3.7054 | 2.1107E-04 | 1.4775E-03 |
| 0.0180 | 0.3171 | 0.0755 | 4.2011 | 2.6600E-05 | 2.1254E-04 |
| 0.0225 | 0.3524 | 0.0403 | 8.7328 | 2.4800E-18 | 4.4700E-17 |
| 0.0270 | 0.3723 | 0.0434 | 8.5837 | 9.1900E-18 | 1.5600E-16 |
| 0.0315 | 0.4175 | 0.0568 | 7.3454 | 2.0500E-13 | 3.0800E-12 |
| 0.0360 | 0.5107 | 0.0397 | 12.8689 | 6.7300E-38 | 1.8900E-36 |
| 0.0405 | 0.5443 | 0.0252 | 21.5674 | 3.6400E-103 | 1.5600E-101 |
| 0.0450 | 0.5039 | 0.0302 | 16.6962 | 1.4000E-62 | 5.3100E-61 |
| 0.0496 | 0.2910 | 0.0331 | 8.8039 | 1.3200E-18 | 2.5100E-17 |
| 0.0541 | 0.1480 | 0.0423 | 3.4988 | 4.6739E-04 | 2.8044E-03 |
| 0.0586 | 0.0640 | 0.0245 | 2.6092 | 9.0761E-03 | 3.6304E-02 |
| 0.0631 | -0.0290 | 0.0472 | -0.6150 | 5.3855E-01 | 5.3855E-01 |

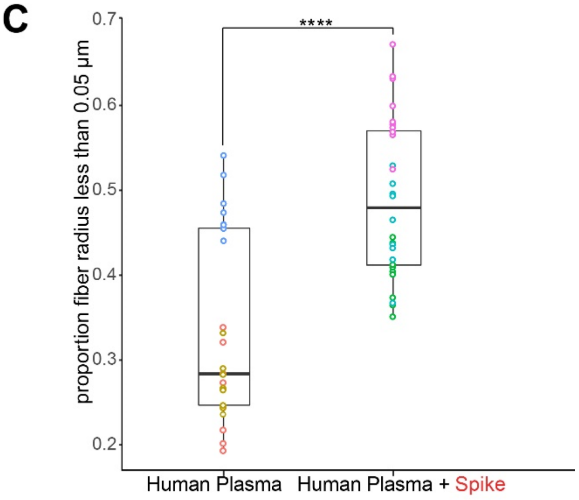


**Fig. S1**. **Analysis of fibrin clot ultrastructure. (A)** SEM images of fibrin clots using human plasma. Top: healthy donor plasma alone. Bottom: healthy donor plasma in presence of Spike protein. 4000X magnification. **(B)** Log odds ratio and corresponding adjusted p values of fibers per measured fiber radius when Spike protein is present compared to plasma alone control. n=3 independent biological replicates. **(C)** Boxplot visualization of the distribution of radii less than 0.05 micron observed in plasma alone compared with plasma in presence of Spike protein. Odds of having observations below 0.05 micron in presence of Spike is 1.92 times (**** $P < 2.2e\text{-}16$) higher than the odds under plasma alone. Generalized linear mixed effect model is used for statistical comparison. $n = 3$ independent biological replicates.

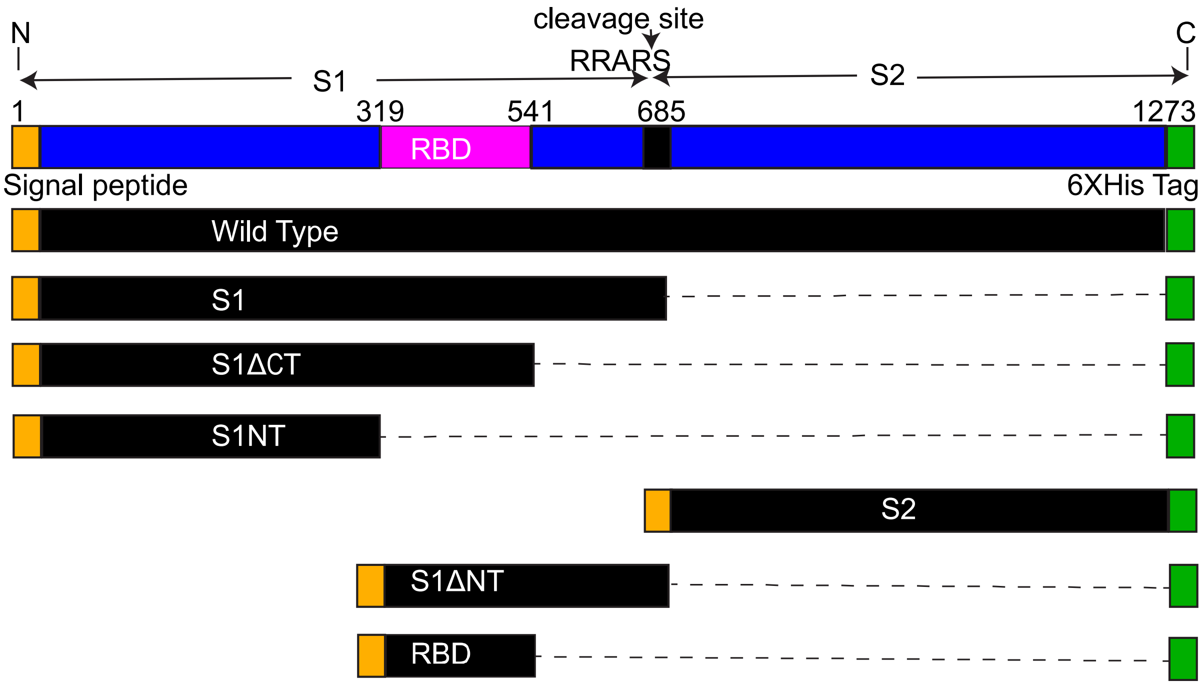


**Fig. S2. Schematic of wildtype full-length Spike and six deletion mutants.** A His6 tag was fused at the C-terminal end of all constructs. S1 C-terminal (S1ΔCT), N-terminal (S1ΔNT), and Receptor binding domain (RBD) (S1NT) deletion mutants are indicated.

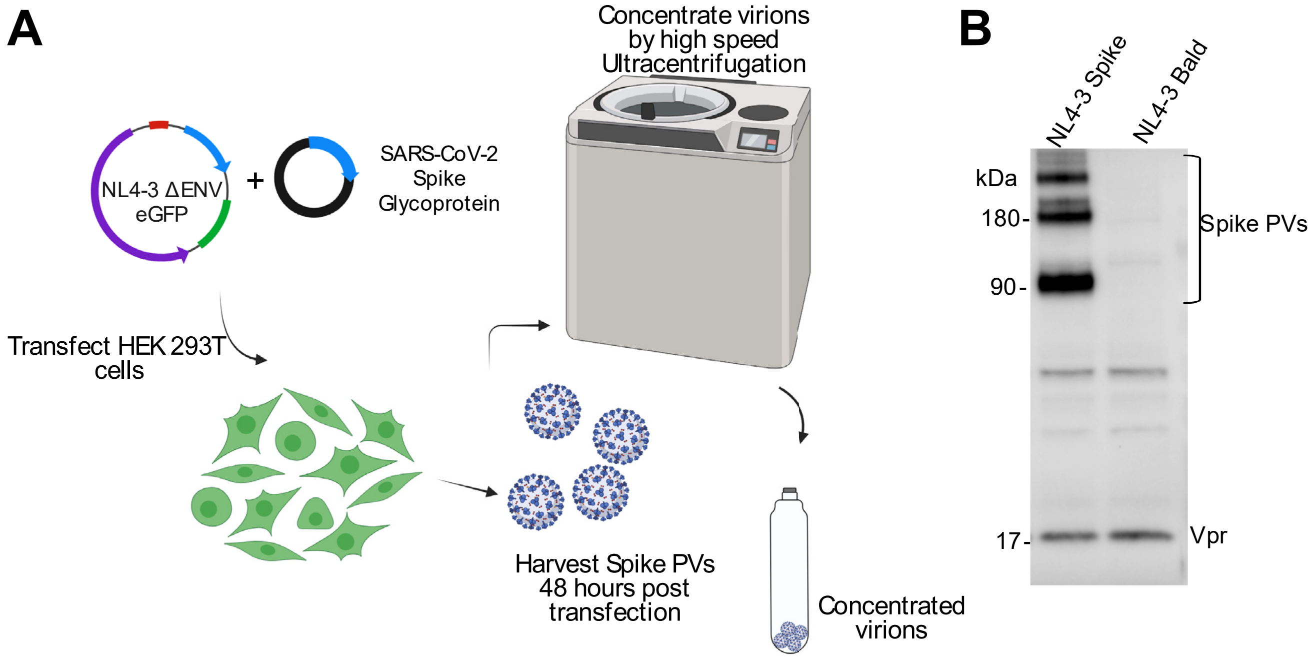


**Fig. S3. Production and validation of PVs. (A)** HIV-1 NL4-3 Δ Env pro-viral DNA vector was co-transfected with the SARS-CoV-2 trimeric Spike glycoprotein expression vector into 293T cells. Forty-eight hours after transfection, supernatant was harvested and Spike PVs were pelleted by ultracentrifugation and collected. **(B)** Western blot of Spike expression of the PVs. NL4-3 Δ Env Spike PVs expressed the S1 and S2 subunits, as well as the cleaved S1 and multimeric forms of the glycoprotein. Comparable levels of HIV Env VPs (Vpr) indicate equivalent expression of the proviral backbone in BALD PVs.

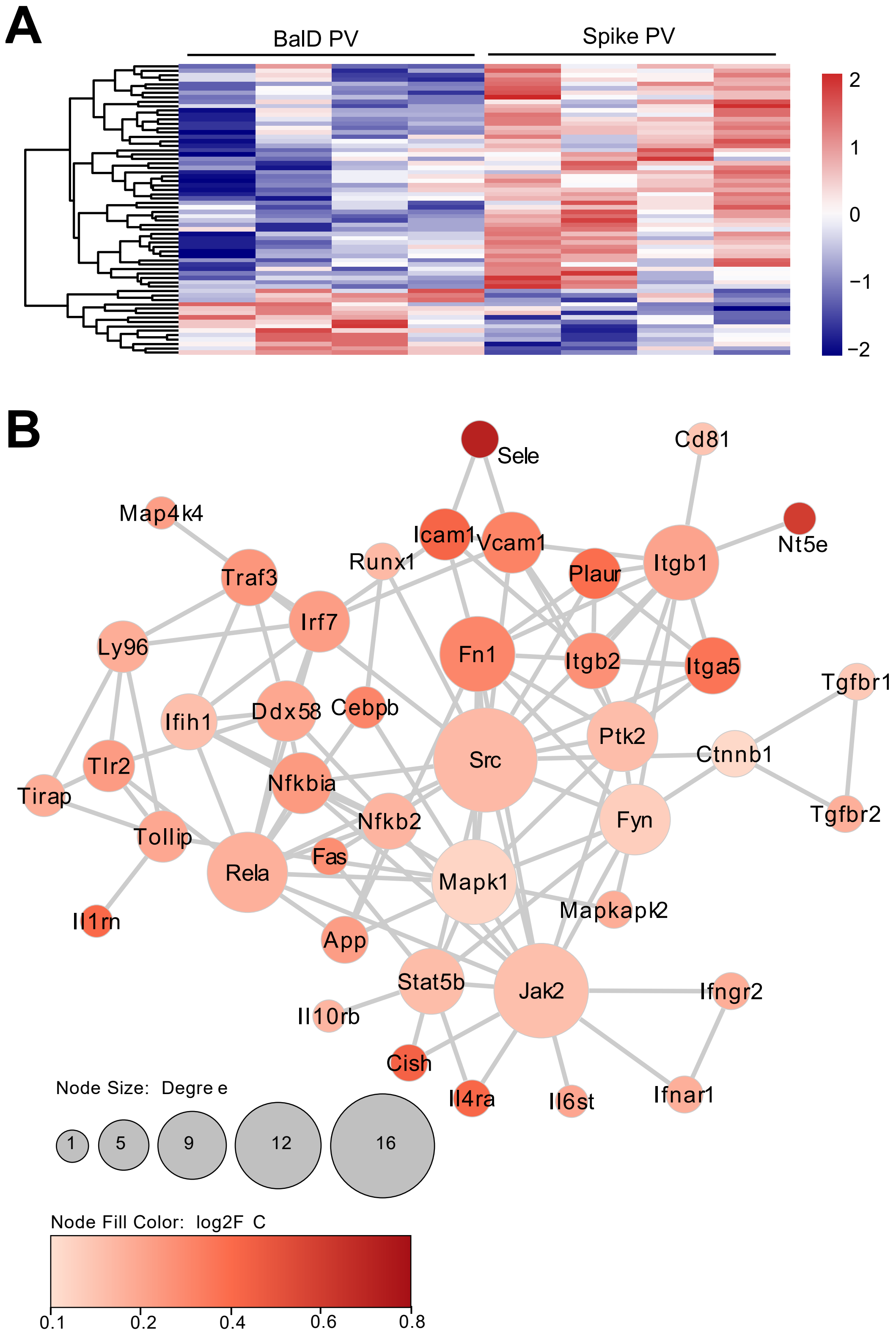
A
BalD PV
Spike PV
2
1
0
−1
−2
B
Sele
Cd81
Map4k4
Nt5e
Icam1
Vcam1
Runx1
Plaur
Itgb1
Traf3
Irf7
Ly96
Fn1
Itgb2
Itga5
Tgfbr1
Ifih1
Ddx58
Cebpb
Ptk2
Src
Ctnnb1
Tlr2
Tirap
Nfkbia
Fyn
Nfkb2
Tgfbr2
Tollip
Fas
Rela
Mapk1
Il1rn
Mapkapk2
App
Stat5b
Jak2
Ifngr2
Il10rb
Cish
Il4ra
Il6st
Ifnar1
Node Size: Degree
1
5
9
12
16
Node Fill Color: log2FC
0.1
0.2
0.4
0.6
0.8

**Fig. S4. SARS-CoV-2 Spike PVs induce an inflammatory gene signature in the lung. (A)** Heatmap of expression of differentially regulated inflammatory genes from the lungs of mice 24 h after injection of Spike PVs or BALD PVs. Data are from 4 biologically independent samples per group. Gene expression is depicted as scaled z score expression (key). **(B)** Largest connected subnetwork of genes significantly upregulated ($P < 0.05$) by Spike PVs based on high-confidence interactions (score > 0.8) from the STRING database: intensity of red shading (key) indicates the extent of the increase in expression after Spike PV injection relative to expression after BALD PV injection; circle size indicates the number of connections per gene in the network.

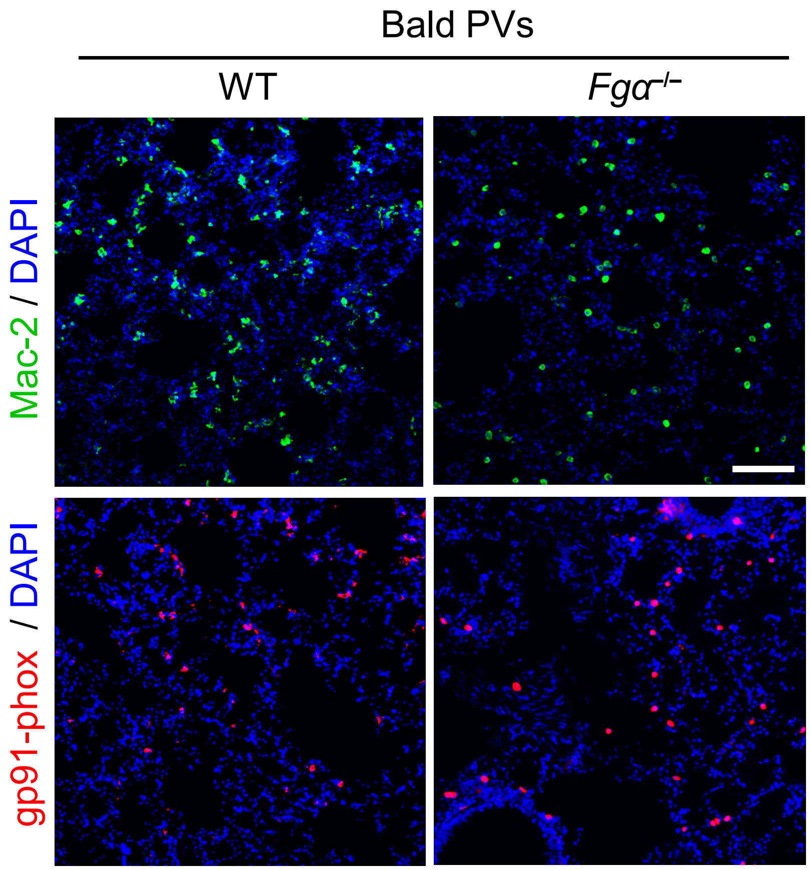


**Fig. S5. Immunohistochemical analysis of lung tissues from WT and *Fga*$^{-/-}$ mice injected with BALD PVs.** Microscopy of lung sections from WT and *Fga*$^{-/-}$ mice 24 h after injection of BALD PVs, showing Mac-2 (green) and gp91-phox (red) immunoreactivity. Scale bar, 50 µm. Representative images from 6 mice per group are shown. Quantification is shown in Fig. 2E.

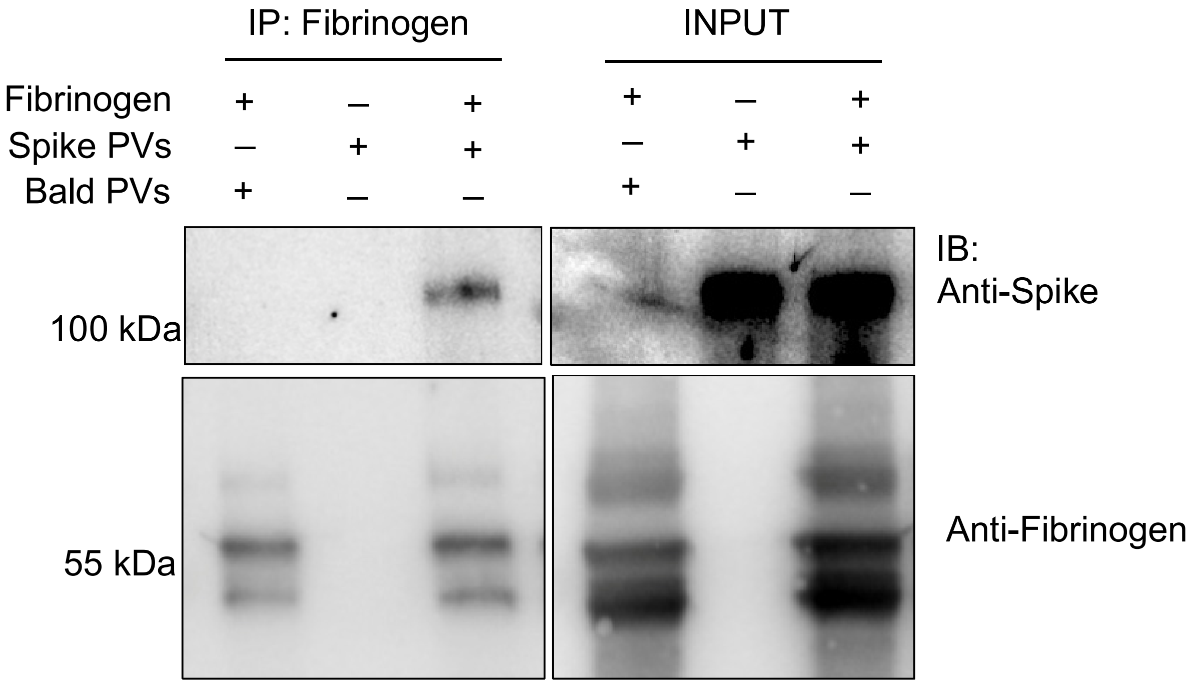


**Fig. S6. Validation of the interaction of Spike PVs with fibrinogen.** Immunoprecipitation of fibrinogen with Spike PVs or BALD PVs, blotted with anti-Spike or anti-fibrinogen. Representative immunoblots from three independent experiments are shown.

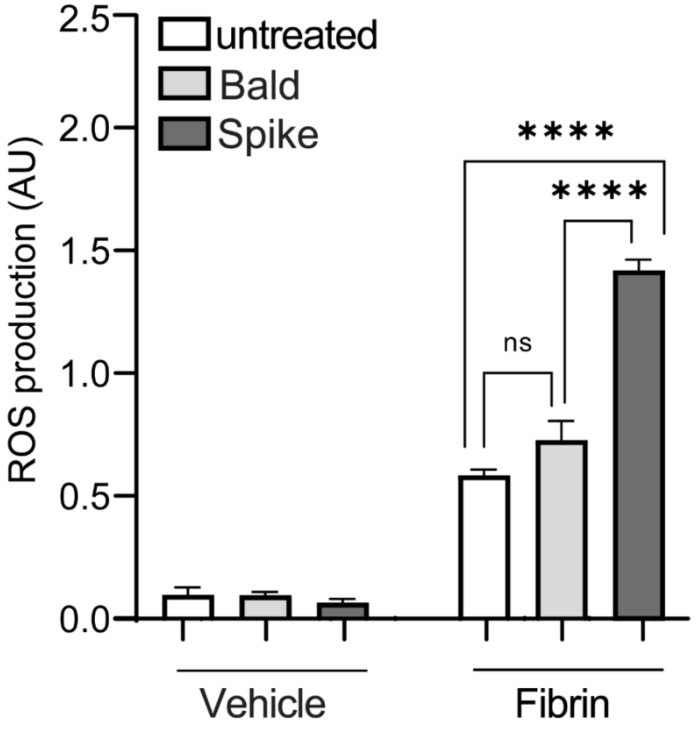


**Fig. S7. Spike PVs enhance fibrin-induced ROS production.** Quantification of ROS production in BMDMs left unstimulated or stimulated for 24 h with fibrin in the presence of BALD PVs or Spike PVs. Values are mean ± s.e.m. of data from three independent experiments. ****$P < 0.0001$ (one-way ANOVA with Tukey's multiple comparisons test). n.s. not significant.

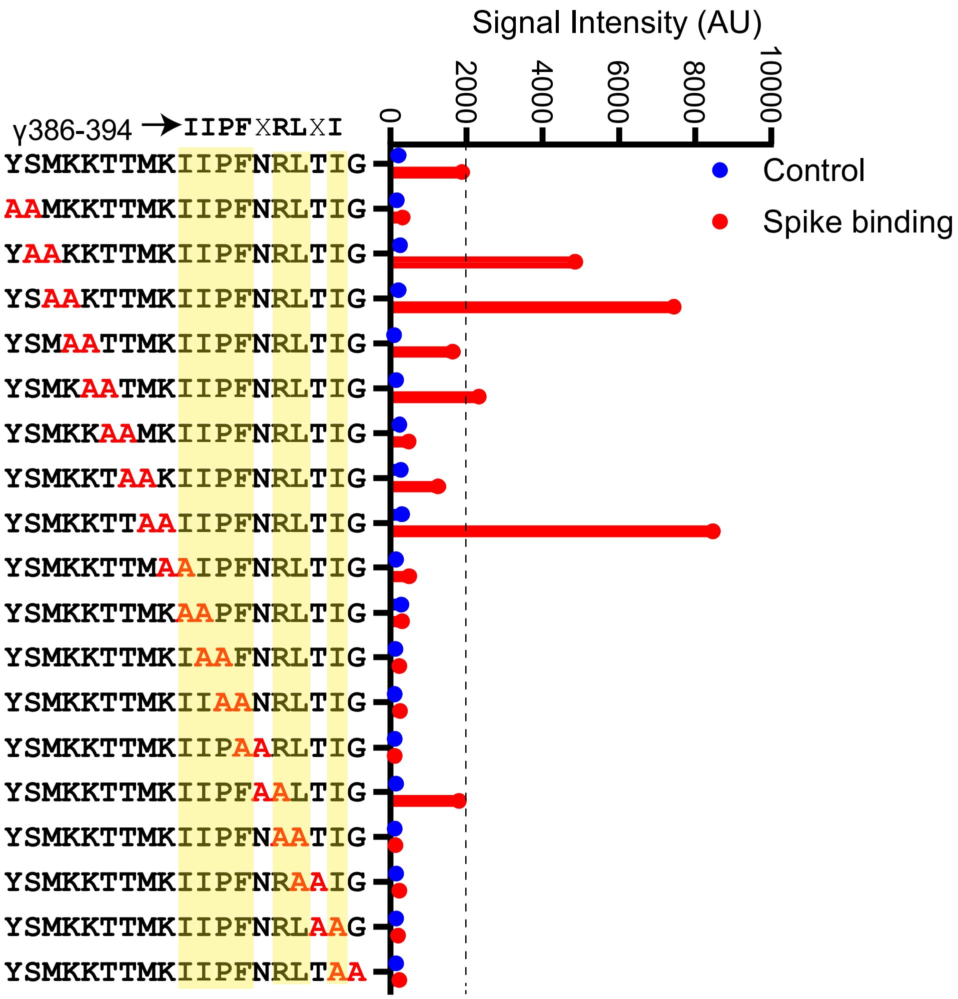


**Fig. S8. Alanine scan mutagenesis peptide array.** Fibrin peptide γ377-395 was subjected to double-alanine scanning mutagenesis and incubated with His-tagged recombinant Spike. Signal intensity bar graph of the binding of Spike to sequential Ala-Ala substituted peptides (red). Control signal is shown in blue. Residues with low signal intensity upon Ala-Ala substitution are required for binding and highlighted in yellow.

**Table S1. Upregulated genes in the lung after SARS-CoV-2 Spike PV administration.** Inflammatory genes differentially upregulated in the lung 24 h after injection of Spike PVs or BALD PVs. Data are from 4 biologically independent samples per group. The adjusted *P* values (adjP) were derived with the Benjamini-Hochberg method to control the false-discovery rate.

| gene | log2FC | p.value | adjP | method | log2FC_thresh |
|---|---|---|---|---|---|
| **Itgb1** | 0.24298757 | 0.00129712 | 0.3986161 | Welch Two Sample t-test | 0 |
| **Mapk1** | 0.08643349 | 0.00218529 | 0.3986161 | Welch Two Sample t-test | 0 |
| **Mapkapk2** | 0.21247061 | 0.00299597 | 0.3986161 | Welch Two Sample t-test | 0 |
| **Cebpb** | 0.33997301 | 0.00358307 | 0.3986161 | Welch Two Sample t-test | 0 |
| **App** | 0.26180232 | 0.00486352 | 0.41570342 | Welch Two Sample t-test | 0 |
| **Il10rb** | 0.1854558 | 0.00643503 | 0.41570342 | Welch Two Sample t-test | 0 |
| **Zbtb7b** | 0.2934328 | 0.00707993 | 0.41570342 | Welch Two Sample t-test | 0 |
| **Fas** | 0.31316965 | 0.00803682 | 0.41570342 | Welch Two Sample t-test | 0 |
| **Tollip** | 0.23036138 | 0.01013979 | 0.41570342 | Welch Two Sample t-test | 0 |
| **Ifnar1** | 0.20485232 | 0.01126423 | 0.41570342 | Welch Two Sample t-test | 0 |
| **Nfkbia** | 0.27687571 | 0.01152716 | 0.41570342 | Welch Two Sample t-test | 0 |
| **Runx1** | 0.17326949 | 0.01176841 | 0.41570342 | Welch Two Sample t-test | 0 |
| **Atg16l1** | 0.17635969 | 0.0139553 | 0.41570342 | Welch Two Sample t-test | 0 |
| **Rela** | 0.20183689 | 0.01482893 | 0.41570342 | Welch Two Sample t-test | 0 |
| **Nox4** | 0.25643446 | 0.01671937 | 0.41570342 | Welch Two Sample t-test | 0 |
| **Ifngr2** | 0.20940353 | 0.01688505 | 0.41570342 | Welch Two Sample t-test | 0 |
| **Il6st** | 0.23597059 | 0.01937659 | 0.41570342 | Welch Two Sample t-test | 0 |
| **Jak2** | 0.15458257 | 0.02111139 | 0.41570342 | Welch Two Sample t-test | 0 |
| **Ube2l3** | 0.13785837 | 0.02180701 | 0.41570342 | Welch Two Sample t-test | 0 |
| **Cd81** | 0.13899363 | 0.02285907 | 0.41570342 | Welch Two Sample t-test | 0 |
| **Npc1** | 0.14615918 | 0.02450686 | 0.41570342 | Welch Two Sample t-test | 0 |
| **Ifih1** | 0.15558846 | 0.02534385 | 0.41570342 | Welch Two Sample t-test | 0 |
| **Il1rn** | 0.43075778 | 0.0263986 | 0.41570342 | Welch Two Sample t-test | 0 |
| **Tlr2** | 0.26977488 | 0.02882483 | 0.41570342 | Welch Two Sample t-test | 0 |
| **Ccbp2** | 0.57589933 | 0.02993537 | 0.41570342 | Welch Two Sample t-test | 0 |
| **Nt5e** | 0.60841058 | 0.03018123 | 0.41570342 | Welch Two Sample t-test | 0 |
| **Tgfbr2** | 0.2068365 | 0.03055609 | 0.41570342 | Welch Two Sample t-test | 0 |
| **Il4ra** | 0.43748981 | 0.03135464 | 0.41570342 | Welch Two Sample t-test | 0 |
| **Tgfbr1** | 0.12396804 | 0.03151048 | 0.41570342 | Welch Two Sample t-test | 0 |
| **Fn1** | 0.33744917 | 0.03235393 | 0.41570342 | Welch Two Sample t-test | 0 |
| **Ddx58** | 0.23203383 | 0.03282107 | 0.41570342 | Welch Two Sample t-test | 0 |
| **Tirap** | 0.2135016 | 0.03351222 | 0.41570342 | Welch Two Sample t-test | 0 |
| **Vcam1** | 0.34917499 | 0.03371891 | 0.41570342 | Welch Two Sample t-test | 0 |
| **Ctnnb1** | 0.07373624 | 0.03375982 | 0.41570342 | Welch Two Sample t-test | 0 |
| **Ly96** | 0.21189091 | 0.03435844 | 0.41570342 | Welch Two Sample t-test | 0 |
| **Sele** | 0.72348825 | 0.03468897 | 0.41570342 | Welch Two Sample t-test | 0 |
| **Traf3** | 0.28478281 | 0.03516047 | 0.41570342 | Welch Two Sample t-test | 0 |
| **Plaur** | 0.41366361 | 0.0370146 | 0.41570342 | Welch Two Sample t-test | 0 |
| **Cish** | 0.45596695 | 0.03831734 | 0.41570342 | Welch Two Sample t-test | 0 |
| **Icam1** | 0.447234 | 0.03843594 | 0.41570342 | Welch Two Sample t-test | 0 |
| **Itga5** | 0.39319074 | 0.03884426 | 0.41570342 | Welch Two Sample t-test | 0 |
| **Fyn** | 0.10849425 | 0.04126186 | 0.41570342 | Welch Two Sample t-test | 0 |
| **Ptk2** | 0.16006092 | 0.04189024 | 0.41570342 | Welch Two Sample t-test | 0 |
| **Irf7** | 0.26311066 | 0.0423173 | 0.41570342 | Welch Two Sample t-test | 0 |
| **Map4k4** | 0.25953665 | 0.04293292 | 0.41570342 | Welch Two Sample t-test | 0 |
| **Nfkb2** | 0.18893622 | 0.04297159 | 0.41570342 | Welch Two Sample t-test | 0 |
| **Itgb2** | 0.30359774 | 0.04493184 | 0.41787499 | Welch Two Sample t-test | 0 |
| **Il1rl2** | 0.20833 | 0.04523157 | 0.41787499 | Welch Two Sample t-test | 0 |
| **Igf2r** | 0.13510123 | 0.0460132 | 0.41787499 | Welch Two Sample t-test | 0 |
| **Src** | 0.17264201 | 0.04821171 | 0.42315816 | Welch Two Sample t-test | 0 |
| **Stat5b** | 0.16193958 | 0.04873669 | 0.42315816 | Welch Two Sample t-test | 0 |

**Table S2. Downregulated genes in the lung after SARS-CoV-2 Spike PV administration.** Inflammatory genes differentially downregulated in the lung 24 h after injection of Spike PVs or BALD PVs. Data are from 4 biologically independent samples per group. The adjusted *P* values (adjP) were derived with the Benjamini-Hochberg method to control the false-discovery rate.

| gene | log2FC | p.value | adjP | method | log2FC_thresh |
|---|---|---|---|---|---|
| H2-Ab1 | -0.24885931 | 0.00128244 | 0.570685761 | Welch Two Sample t-test | 0 |
| Cd209g | -0.83051076 | 0.008923569 | 0.998702878 | Welch Two Sample t-test | 0 |
| H2-Aa | -0.25233333 | 0.018505218 | 0.998702878 | Welch Two Sample t-test | 0 |
| Ccl11 | -0.37658203 | 0.021602196 | 0.998702878 | Welch Two Sample t-test | 0 |
| Slamf7 | -0.24931128 | 0.024678967 | 0.998702878 | Welch Two Sample t-test | 0 |
| Casp1 | -0.15752512 | 0.026866216 | 0.998702878 | Welch Two Sample t-test | 0 |
| Ccr6 | -0.85150502 | 0.028325191 | 0.998702878 | Welch Two Sample t-test | 0 |
| H2-DMa | -0.24780279 | 0.029033537 | 0.998702878 | Welch Two Sample t-test | 0 |
| Hcst | -0.42482418 | 0.029282175 | 0.998702878 | Welch Two Sample t-test | 0 |
| Il11ra1 | -0.5749819 | 0.032494474 | 0.998702878 | Welch Two Sample t-test | 0 |
| Cd74 | -0.14564625 | 0.035484511 | 0.998702878 | Welch Two Sample t-test | 0 |
| Cd55 | -0.17085985 | 0.036489801 | 0.998702878 | Welch Two Sample t-test | 0 |
| H2-DMb2 | -0.51091377 | 0.040944006 | 0.998702878 | Welch Two Sample t-test | 0 |
| Gpr183 | -0.32944692 | 0.042115214 | 0.998702878 | Welch Two Sample t-test | 0 |
| Cd79b | -0.64084215 | 0.042690857 | 0.998702878 | Welch Two Sample t-test | 0 |

**Table S3. Characteristics of COVID-19 and control patients included in the fibrin autoantibody screen.** SD: standard deviation.

| Patient Characteristics | Severe COVID-19 n=19 | Mild COVID-19 n=20 | Asymptomatic COVID-19 n=15 | Healthy Control n=9 | Non-COVID Respiratory Illness n=8 |
|---|---|---|---|---|---|
| **Age, y** | | | | | |
| Median | 60 | 44 | 53 | 37 | 61 |
| Mean (SD) | 60.05 (13.9) | 47.74 (12.6) | 54.9 (14.6) | 40.4 (9) | 64.9 (15.8) |
| **Sex** | | | | | |
| Male | 16 | 14 | 8 | 4 | 8 |
| Female | 3 | 6 | 7 | 5 | 0 |
| **Level of Care** | | | | | |
| Mechanical ventilation | 19 | 0 | 0 | 0 | – |
| **Mortality** | | | | | |
| | 3 | 0 | 0 | 0 | – |
| **Medical History** | | | | | |
| Hypertension | 14 | 7 | 6 | 0 | – |
| Type 2 diabetes | 10 | 2 | 4 | 0 | – |
| Obesity | 8 | 3 | 3 | 0 | – |
| Coronary artery disease | 1 | 1 | 0 | 0 | – |
| Congestive heart failure | 2 | 1 | 1 | 0 | – |
| Stroke | 1 | 0 | 0 | 0 | – |
| Chronic kidney disease | 1 | 0 | 1 | 0 | – |
| Chronic liver disease | 0 | 0 | 1 | 0 | – |
| COPD | 1 | 0 | 1 | 0 | – |
| Asthma | 0 | 0 | 1 | 0 | – |
| Organ transplant | 0 | 1 | 0 | 0 | – |
| Immunocompromised | 0 | 1 | 0 | 0 | – |
| HIV | 0 | 0 | 1 | 0 | – |
| None | 3 | 8 | 4 | 0 | – |
| **Symptoms** | | | | | |
| Fever | 13 | 13 | 0 | 0 | – |
| Cough | 14 | 8 | 0 | 0 | – |
| Shortness of breath | 15 | 5 | 0 | 0 | – |
| Sore throat | 3 | 5 | 0 | 0 | – |
| Chest pain | 2 | 3 | 0 | 0 | – |
| Loss of taste/smell | 0 | 1 | 0 | 0 | – |
| Headache | 3 | 4 | 0 | 0 | – |
| Myalgia | 4 | 6 | 0 | 0 | – |
| Chills | 3 | 6 | 0 | 0 | – |
| Diarrhea | 3 | 3 | 0 | 0 | – |

**Movie S1. Confocal imaging of z-stacks of Spike and fibrinogen in the lung.** High-resolution confocal imaging of Spike and fibrinogen in fixed mouse lung tissue after SARS-CoV-2 Spike PV administration. z-stacks were acquired (30 µm in 1-micron steps) with FluoView on the Olympus BX61W1 confocal microscope with a 20× Olympus objective.

**Movie S2. Confocal imaging of Spike and fibrinogen in the lung.** Confocal 3D volume projections of lung after SARS-CoV-2 Spike PV administration showing fibrinogen (red) and Spike (green) overlapping around vessels (yellow).
